# Explosive Synchronization-Based Brain Modulation Reduces Hypersensitivity in The Brain Network: A Computational Model Study

**DOI:** 10.1101/2021.10.18.464748

**Authors:** MinKyung Kim, Richard E. Harris, Alexandre F. DaSilva, UnCheol Lee

## Abstract

Fibromyalgia (FM) is a chronic pain condition that is characterized by hypersensitivity to multi-modal sensory stimuli, widespread pain, and fatigue. We have previously proposed explosive synchronization (ES), a phenomenon wherein a small perturbation to a network can lead to an abrupt state transition, as a potential mechanism of the hypersensitive FM brain. Therefore, we hypothesized that converting a brain network from ES to general synchronization (GS) may reduce the hypersensitivity of FM brain. To find an effective brain network modulation to convert ES into GS, we constructed a large-scale brain network model near criticality (i.e., an optimally balanced state between order and disorders), which reflects brain dynamics in conscious wakefulness, and adjusted two parameters: local structural connectivity and signal randomness of target brain regions. The network sensitivity to global stimuli was compared between the brain networks before and after the modulation. We found that only increasing the local connectivity of hubs (nodes with intense connections) changes ES to GS, reducing the sensitivity, whereas other types of modulation such as decreasing local connectivity, increasing and decreasing signal randomness are not effective. This study would help to develop a network mechanism-based brain modulation method to reduce the hypersensitivity in FM.

**Author summary:** Phase transitions, the physical processes of transition between system states in nature, are divided into two broad categories: first and second-order phase transitions. For example, boiling water presents abrupt transition (a first-order) along with high sensitivity to temperature change, distinct from gradual magnetization near Curie temperature (a second-order). Recently, we found that chronic pain shows specific brain network configurations that can induce the first-order transition, so-called ‘explosive synchronization.’ In this modeling study, we tried to identify a modulation method that can convert a first-order transition into a second-order transition in the brain network, expecting that it may inhibit the hypersensitivity in chronic pain. We found that increasing structural connectivity of hubs changes the type of phase transition in the brain network, significantly reducing network sensitivity.

## Introduction

Hypersensitive responses to external stimuli have been widely observed in various physical and biological systems such as cascading failures in power-grids, abrupt state transitions in an electronic circuit, abrupt loss and recovery of consciousness in general anesthesia, and epileptic seizures in the brain (Boccaletti et al., 2016; Chen et al., 2013; C.-Q. Wang et al., 2017). Fibromyalgia (FM), a chronic pain disorder, is characterized by fatigue, poor memory, sleep problems, and mood disturbance (Hawkins, 2013; Menzies, 2016). Many of these individuals also present a hypersensitive response to external sensory stimuli, which is regarded to involve central sensitization associated with structural and functional changes in the brain (Harris et al., 2013; Harte et al., 2018). In our previous study, we found that explosive synchronization (ES; a 1^st^ order phase transition in a network) to be an underlying mechanism of the hypersensitivity in FM brain (U. Lee et al., 2018). A strong positive correlation was detected between the strength of ES conditions and chronic pain intensity in FM patients. This suggests that specific topological and functional network configurations of the brain could induce the 1^st^ order phase transition (abrupt state transition) against an external stimulus (M. Kim et al., 2016, 2017; U. Lee et al., 2018).

Global synchronization in a network is initiated by the merging of synchronization clusters across nodes. In a brain network following a general synchronization (GS) path, hubs (nodes with intense connections) lead a global synchronization of the network, forming a large synchronization cluster (Gómez-Gardeñes et al., 2007). Once a large synchronization cluster has formed, it grows gradually, absorbing smaller local clusters. However, if the growth of the largest cluster is somehow suppressed by pharmacological, neurological, or pathological types of perturbation, it allows disconnected clusters to grow until the network reaches a critical threshold wherein a small stimulus triggers a singular explosive unification of all clusters termed explosive synchronization (ES). In these synchronization processes, hubs play a central role in creating a giant synchronization cluster in the brain network (M. Kim et al., 2017; Zhang et al., 2015). Therefore, we expected modulating the hubs in the brain network can be the most effective and practical stimulation target to convert the type of state transition, from ES to GS, in brain networks.

A previous model study demonstrated the possibility to convert an ES network into a GS network by modulating the ratio of inhibitory to excitatory connectivity (Zhang et al., 2016). However, modulating the ratio of inhibitory connectivity in the global brain network may not be feasible with current stimulation techniques. Furthermore, it has not been tested if the brain network state is near criticality, which presents complex brain dynamics of a conscious state, is convertible from ES into GS by a brain modulation. In principle, reversing the core mechanism of ES may convert an ES network into a GS network, but this would require reversing suppression of the giant synchronization cluster formation. In essence, this may change an abrupt state transition to a gradual state transition as well as significantly reduce the network sensitivity. We hypothesized that enhancing the hub connectivity may be a way to facilitate the giant synchronization cluster formation, which in turn can convert an ES network to one of GS.

To justify the hypothesis, we constructed a large-scale human brain network model occurring ES, using a modified Stuart-Landau oscillator model on the anatomically informed human brain network structure (H. Kim et al., 2018). After constructing ES brain network models, two different network properties were modulated as control parameters for the brain network modulation: (1) the local structural connectivity of brain regions and (2) the randomness of node dynamics within a certain diameter centered on a target node. The correlation between node degrees and frequencies, which is one of the ES conditions, was measured at critical states of the brain networks before and after the modulation. To evaluate which is the most effective type of brain network modulation for reducing the brain network sensitivity, we directly applied an external stimulus to the brain network models and measured the change of brain network sensitivity with surrogate measures— responsivity and Lempel Ziv complexity (M. Kim & Lee, 2020). The schematic diagram of the study design is illustrated in Figure 1.

**Figure 1.**
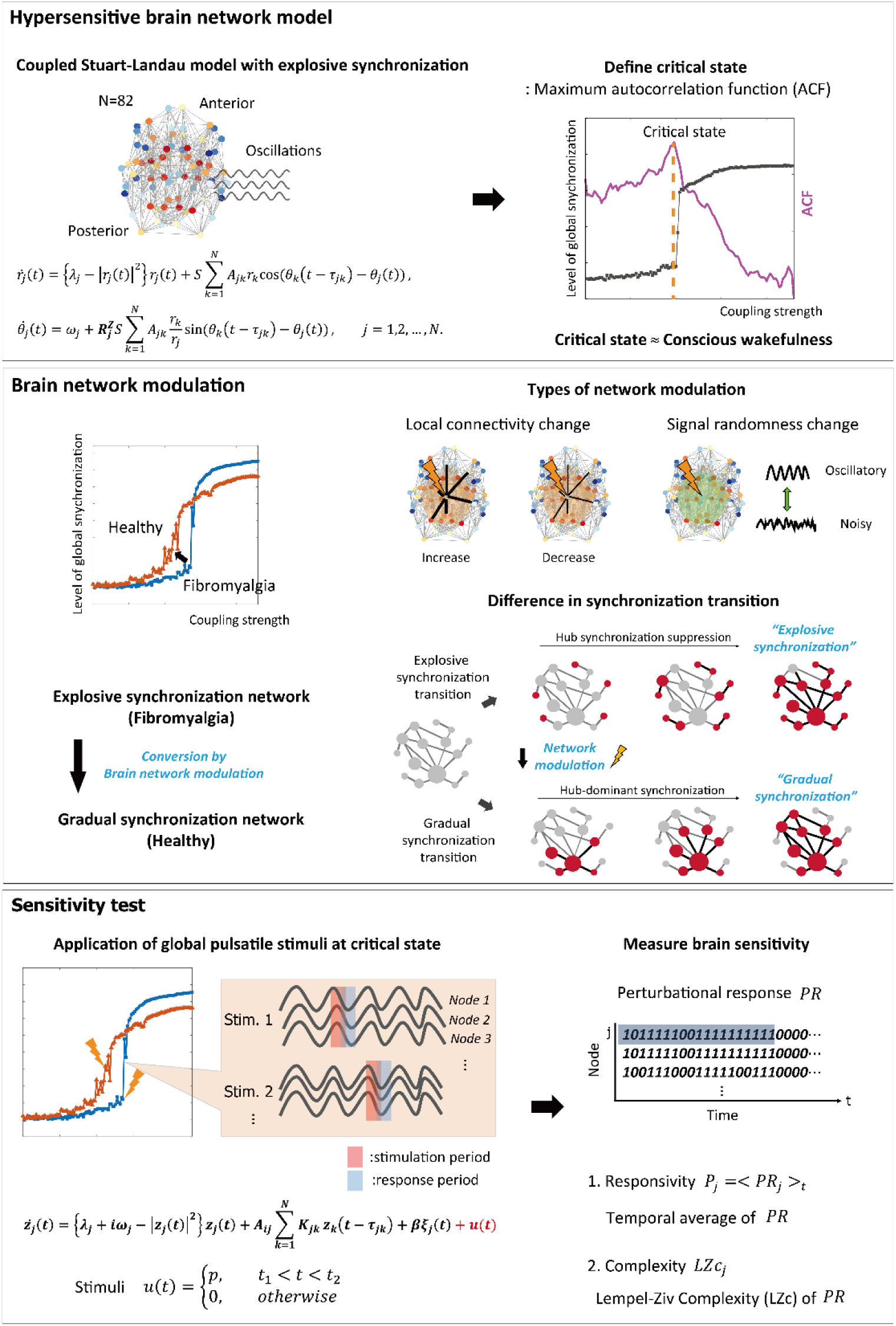
Schematic diagram of the study design. We simulated the fibromyalgia (FM) brain network using a model with the explosive synchronization (ES) mechanism. The ES brain network model was constructed by using a modified coupled Stuart-Landau oscillator on an anatomically informed human brain network structure. We found a critical state of ES brain network model by calculating autocorrelation function (ACF) to simulate the FM brain during conscious wakefulness. Four different types of network modulation (local structural connectivity increase and decrease, signal randomness increase and decrease) with thirty different target brain regions were applied to the model to investigate which modulation types and which target brain regions can convert an ES network to one of general synchronization (GS) thereby reducing the sensitivity. Then we induced external stimuli to the brain networks at a critical state before and after the modulation, evaluated sensitivity (responsivity and complexity) of the brain network responses, and compared the sensitivity between the networks before and after the modulation.

## Results

### Increasing regional brain connectivity converts abrupt transition into gradual transition

Four types of brain network modulation (local structural connectivity increase: CI; local structural connectivity decrease: CD; local randomness (bifurcation parameter) increase: RI; local randomness (bifurcation parameter) decrease: RD) were tested to convert an ES network into a GS network. For each type of modulation, we applied the modulation to the brain networks centered on thirty different highest-degree nodes to investigate the effect of regional modulation. Figure 2 presents representative examples for the shape of synchronization transition before (blue line) and after (red line) the modulation. The examples are the shapes of synchronization transition of brain network modulation targeting the right precuneus for each type of modulation. The simulated brain network before the modulation showed an abrupt synchronization transition near the critical state (blue lines in Figure 2A-2D), which reflects ES (the 1^st^ order phase transition). Only CI modulation converted the type of synchronization transition from ES to GS (red line in Figure 2A), whereas the other types of modulation (CD, RI, and RD) did not (red lines in Figure 2B,2C, and 2D). The shapes of synchronization transition targeting other nodes of all types of modulation were presented in supplementary materials (Supporting Information Figure S1-S4). The CI modulation targeting other nodes mostly showed the shape change of synchronization transition from abrupt to gradual (Supporting Information Figure S1). Other types of modulation (CD, RI, and RD) did not show the shape change in most of the nodes (Supporting Information Figure S2-S4).

**Figure 2.**
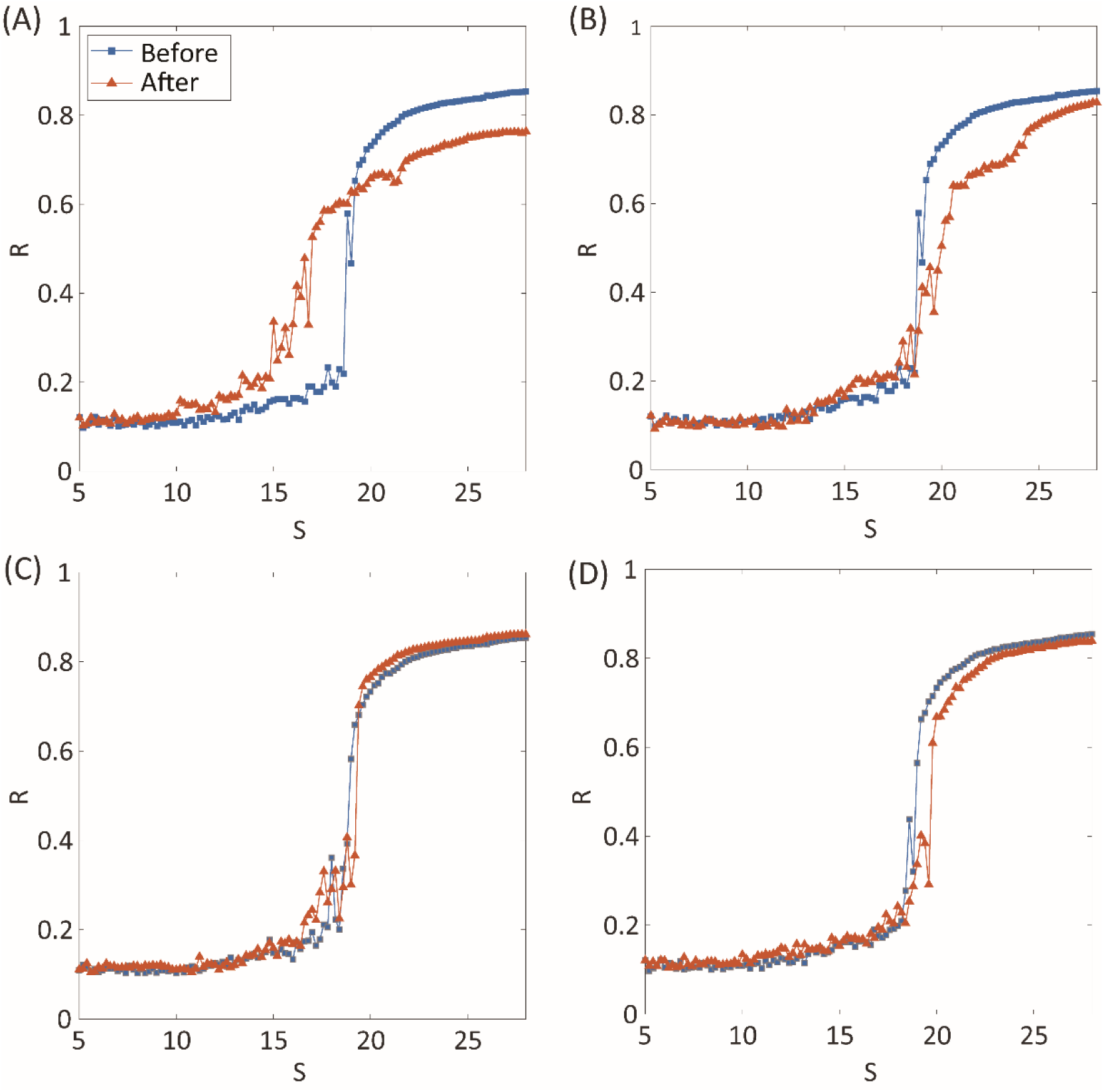
Synchronization transition shapes of four types of network modulation: (A) local structural connectivity increase: CI; (B) connectivity decrease: CD; (C) signal randomness (bifurcation parameter) increase: RI; (D) signal randomness (bifurcation parameter) decrease: RD. For each type of modulation, thirty different brain regions were targeted and modulated respectively. The node modulation centered around the left precuneus is shown as an example of node modulation. A blue (orange) line indicates levels of network synchronization *R* along with a change of the coupling strength *S* of the model before (after) modulation. Only CI shows relatively gradual synchronization in the network after the modulation. The abrupt transition near the critical point, which is one of the characteristics of ES, is relatively maintained for CD, RI, and RD modulation.

### Increasing regional brain connectivity mitigates ES condition

We examined a correlation between node degree and node frequency (*ρ*^*deg*−*freq*^) in the brain network to investigate whether the four types of modulation disrupt a typical ES condition. In generic network models, a larger positive *ρ*^*deg*−*freq*^ induces ES with a higher probability. Figure 3A-3D respectively show the *ρ*^*deg*−*freq*^ values before and after the four types of modulation for the thirty target nodes. The gray shaded areas in Figure 3 indicate the mean ± standard error of *ρ*^*deg*−*freq*^ before the modulation and the square dots with error bars indicate the mean ± standard error of *ρ*^*deg*−*freq*^ after the modulation. The name of the target brain regions is presented on the x-axis. We found that only CI modulation gives rise to significant decreases of *ρ*^*deg*−*freq*^ for the most target node modulations (Left pallidum, thalamus, hippocampus, amygdala, temporal pole, insula, entorhinal cortex, precentral cortex, postcentral cortex, isthmus cingulate, right thalamus, temporal pole, cortex, right precuneus, entorhinal cortex, and accumbens area; t-test, *p<0.05, Figure 3A), while the CD modulation showed an increase of *ρ*^*deg*−*freq*^ for some target nodes (Left and right putamen, right amygdala, left and right pallidum, right thalamus, left amygdala, left and right caudate, right insula, and right entorhinal cortex, Figure 3B). The RI and RD modulations did not change *ρ*^*deg*−*freq*^ or only a few target node modulations induced alternation (Figure 3C, and 3D). The comparison among *ρ*^*deg*−*freq*^ values for four types of modulation is presented in Supporting Information Figure S5. The CI modulation was significantly lower than other types of modulation. In addition, we investigated the change of signal properties such as a global synchronization and a signal amplitude, which is associated with the responsiveness of a network against stimulation (M. Kim & Lee, 2020). Only CI modulation significantly increased the global synchronization and the amplitude of the signal (Supporting Information Figure S6), which implies the decreased responsiveness and the mitigated ES condition.

**Figure 3.**
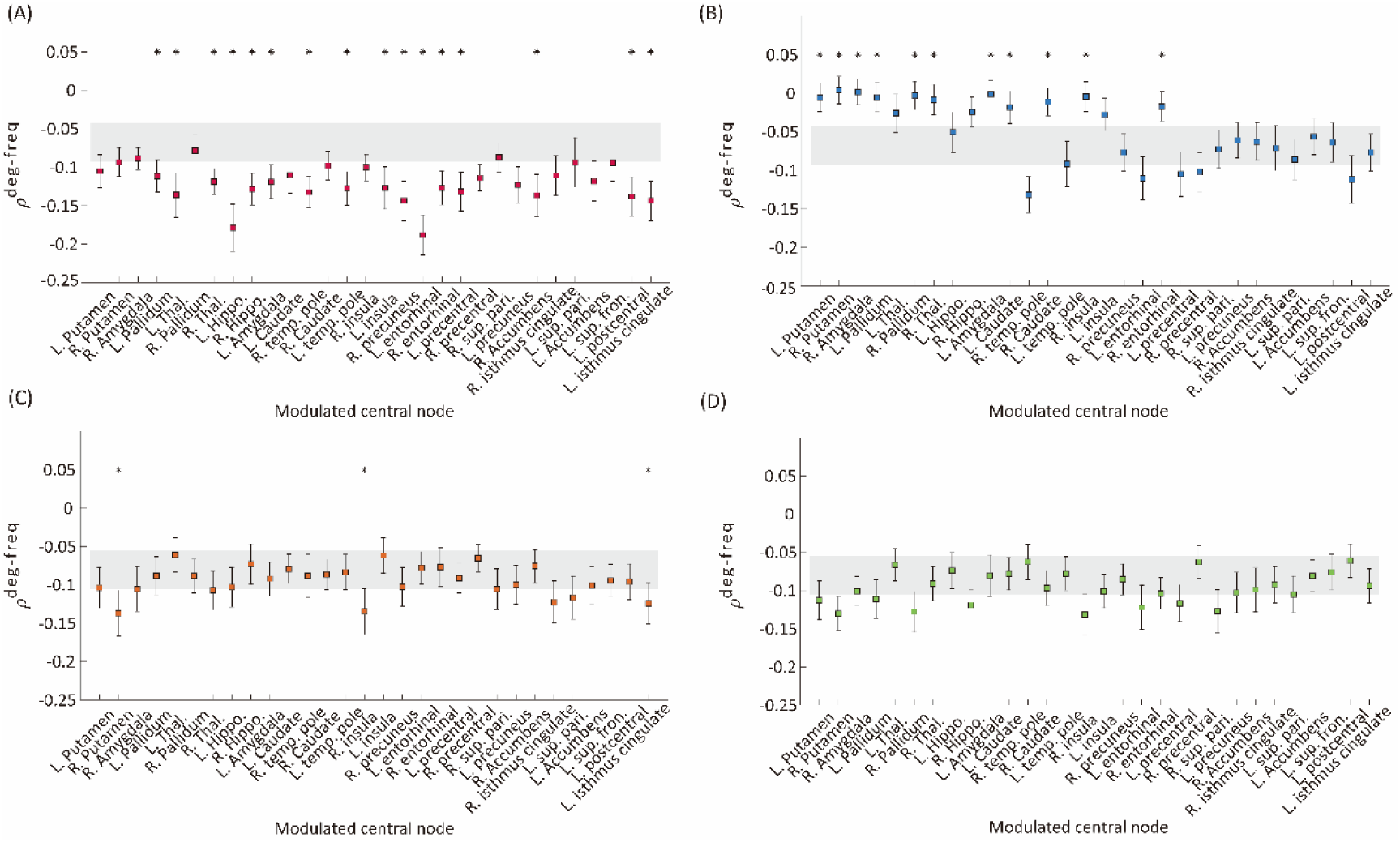
Correlation values between node degree and frequency *ρ*^*deg*−*freq*^ of the networks before and after the four types of modulation: (A) CI, (B) CD, (C) RI, and (D) RD. The grey area indicates the mean±standard errors of *ρ*^*deg*−*freq*^ over 30 different initial conditions of brain networks before the modulation. The colored square with error bars indicates the mean±standard errors of *ρ*^*deg*−*freq*^ over 30 different initial conditions of brain networks after the modulation. Each marker indicates the centered target node for the modulation. A target node presenting a significant change after the modulation is marked with ‘*’ (t-test, *p<0.05). The CI modulation induces a significant decrease of *ρ*^*deg*−*freq*^ for most of the target nodes. In contrast, other types of modulation induce significant changes for the much smaller number of nodes (CD and RI). The RD modulation shows no change for all nodes.

### Increasing regional brain connectivity decreases network sensitivity

Next, we implemented an external stimulus and compared the changes in brain network responses before and after the modulation to directly evaluate the brain network sensitivity change. Responsivity *P* and complexity *LZc* were calculated to evaluate the brain network sensitivity with perturbed signals during 300 msec after the stimulation. Only CI modulation with specific target brain regions showed decreases in both responsivity and complexity. Figure 4 shows the responsivity (left) and complexity (right) for the CI modulation. The names of the target brain regions are presented in the x-axis. The gray area covers the 25% to 75% values of *P* and *LZc* before the CI modulation. The blue and green shaded areas in Figure 4A and 4B respectively denote the 25% to 75% quantiles of *P* and *LZc* after the CI modulation for 30 different target brain regions. The red star markers indicate the effective target brain regions where the network sensitivity is significantly decreased (t-test, *p<0.05). Increasing the regional brain connectivity centered on subcortical regions and some cortical hub regions (Left putamen, pallidum, amygdala, insula, entorhinal cortex, accumbens, isthmus cingulate cortex, right putamen, amygdala, pallidum, hippocampus, temporal pole, caudate nucleus, insula, precuneus, and entorhinal cortex) significantly decrease the brain network sensitivity. The responsivity of the brain networks in the other types of modulation (RI and RD) was also decreased significantly, however, the changes in responsivity were not (Supporting Information Figure S7).

**Figure 4.**
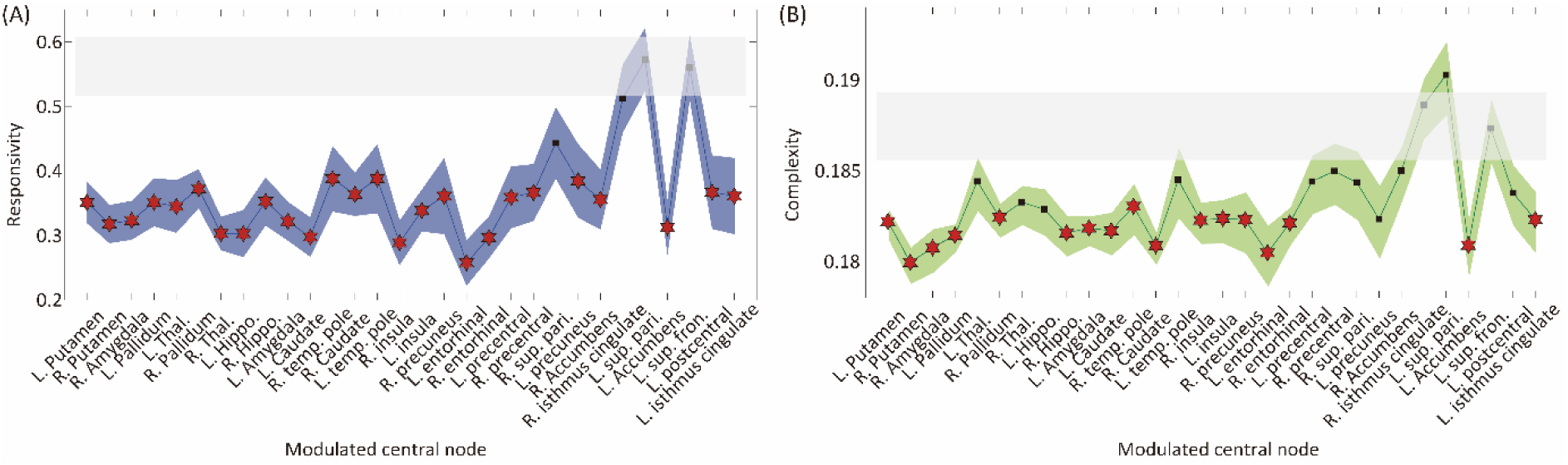
Changes in brain network responsivity and complexity after the CI modulation. The (A) responsivity and (B) complexity of the brain networks before and after the CI modulation are presented. The grey area covers 25% to 75% values of network responsivity (sensitivity) before the CI modulation over different initial conditions. The blue (green) colored area covers 25% to 75% values of network responsivity (complexity) after modulation over different initial conditions. A target node presenting significantly different responsivity (complexity) is marked as ‘*’ (t-test, *p<0.01). The CI modulation to the right and left insula, right precuneus, and left isthmus cingulate cortex results in decreased responsivity and complexity in the brain network.

### Hub nodes are effective target sites to reduce brain network sensitivity

We hypothesized that the hub nodes in the brain network can be effective target sites to reduce brain sensitivity. Therefore, we evaluated the relationship between the node degree of target brain regions and the sensitivity of the node modulation. Figure 5A (5B) presents the relationship between the average node degree of target brain regions and the responsivity (complexity) of the target node CI modulation. Both responsivity and complexity showed significant negative Spearman correlations with the node degree of target brain regions (−0.52 and −0.43, respectively), suggesting that the CI modulation targeting hub nodes gives rise to lower responsivity and complexity than targeting peripheral nodes. The large correlation between network sensitivity and node degree implies the existence of effective target sites in the brain network, which also reveals the possibility of developing a systematic brain modulation method with better outcomes considering the network topology.

**Figure 5.**
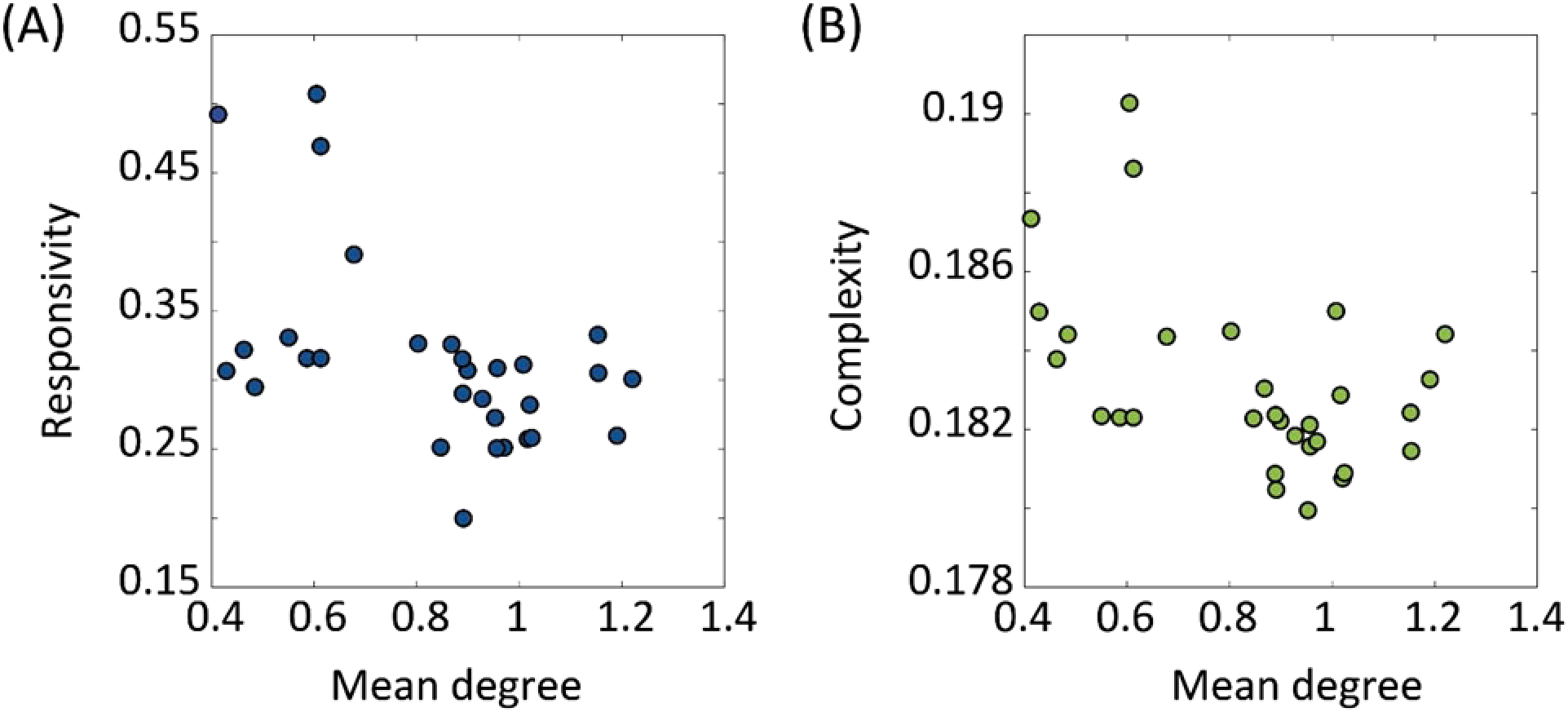
The relationship between the node degree of target brain regions and the sensitivity after the CI modulation. The sensitivity was evaluated by responsivity and complexity. The relationships between the node degree of target brain regions and the (A) responsivity and (B) complexity are presented. The Spearman correlation coefficient between the network responsivity (complexity) and the average degree of target nodes is −0.52 (−0.43). The CI modulation to larger degree nodes (i.e. hubs) induces more decrease in the brain network sensitivity.

## Discussion

Hypersensitivity of ES may provide an opportunity for a system to have flexible adaptation to external stimuli with high susceptibility and fast response, but also it implies high risks of unwanted excessive responses that might cause a neurologic problem in the brain which may explain previous neuroimaging findings in FM. Empirical evidence of ES in FM brain network and the possibility for converting ES into GS network in previous model studies motivated us to investigate brain modulation methods that could reduce hypersensitivity of the brain network at the fundamental level. We found that increasing the regional brain connectivity, especially centered on hubs, changed the ES brain network to GS brain network, disrupted the ES condition, and reduced network sensitivity against an external stimulus. Considering a central role of hubs in FM (Kaplan et al., 2019) and giant synchronization cluster formation, changing hub connectivity in the brain network could potentially reverse the key mechanism of ES, facilitating giant synchronization cluster formation in the network, so that in principle the network is converted from ES (a 1^st^ order phase transition) into GS (a 2^nd^ order phase transition) with reduced brain network sensitivity.

### Converting ES into GS in the brain network

Previous computational modeling studies have identified network conditions to prevent the onset of ES (Dai et al., 2020; Zhang et al., 2016). The authors have found that ES could be changed to GS if the population of inhibitory connections was larger than a threshold (about 10%) in the networks consisting of inhibitory and excitatory connections. The ratio of the inhibitory connections required for converting ES into GS depended on the network size (the total number of nodes) and the average node degree (the average number of connections among nodes). Such existence of the threshold implies that converting ES into GS is controllable by modulating the population of inhibitory connections. However, it is difficult to directly apply this method to a large-scale human brain network due to its complex hierarchical modular structure and complex dynamics. In addition, it has not been tested in network dynamics near a critical state, presumably reflecting brain network dynamics in a conscious state. Therefore, we instead searched a practical way to convert ES into GS in the brain network, modulating local brain connectivity and local brain signal randomness that can facilitate a giant synchronization cluster formation, that is, reversing the core mechanism of ES. In addition, targeting local brain regions with intense connectivity such as hubs in the brain network is feasible with current non-invasive brain stimulation tools.

### Hub as an effective target site

We selected hubs in the brain network as the most promising target site for the brain network modulation because brain network hubs are the most powerful candidate that can control the whole brain network with local modulation. Hubs have been known that they show the largest network controllability in the model (Gu et al., 2015) and play a central role in chronic pain (Kaplan et al., 2019). In addition, hub structure in the brain network plays as a major determinant in global synchronization and desynchronization (Schmidt et al., 2015; Vlasov & Bifone, 2017). Topologically, the hierarchical hub structure in the brain network called “rich club,” is a highly connected and centralized collection of nodes that occupies only 10% of the brain network but facilitates 70% of the communication pathways across the brain regions (Schmidt et al., 2015; van den Heuvel & Sporns, 2011), which can be one of the reasons why hubs are the most effective target sites to promote or suppress global synchronization in the brain network.

Recent brain imaging studies have presented that the hub or the membership of the rich club is varied with the intensity of clinical pain. The posterior insula, primary somatosensory, and motor cortices (S1/M1) belonged to the rich club only in FM patients with the highest clinical pain intensity, which was not observed in patients with low pain intensity and healthy controls. The altered hub topology suggests that pronociceptive regions such as insula appear to have acquired new hub status that would influence the quality and quantity of information processing in the brain network (Kaplan et al., 2019). Moreover, the altered hub topology with new hubs may enhance heterogeneity of the brain network, suppressing the onset of global synchronization and changing the brain network closer to the ES network. Therefore, we propose that generating a dominant hub node whose local connectivity is enhanced via CI modulation can successfully convert ES to GS. The enhanced hub connectivity can lead to a gradual synchronization propagation starting from the dominant hub node to other nodes, instead of a sudden singular unification near a critical point.

### Increasing local structural connectivity in brain network

In this study, we showed that only increasing local structural connectivity of target nodes can mitigate the ES condition with a more negative correlation between node degree and frequency (Figure 3). A positive correlation between node degree and frequency is a well-known network configuration for ES, suppressing a giant synchronization cluster formation (Gómez-Gardeñes et al., 2011). If higher frequencies were distributed to hubs in a network (i.e. a positive correlation between node degree and frequency), hubs would suppress the overall network synchronization until the network reaches a critical point because higher frequency oscillators have small/irregular amplitudes and incoherent phases in a network (Gollo et al., 2015; Honey et al., 2007; Moon et al., 2015, 2017). On the contrary, if slower frequencies were distributed to hubs (i.e. a negative correlation between node degree and frequency), hubs would tend to increase the local synchronization around them and progressively expand the synchronization throughout the whole network (Moon et al., 2015).

We evaluated how the network sensitivity is significantly changed by a direct brain stimulus after the four types of brain network modulation (increasing/decreasing brain regional connectivity around target nodes and increasing/decreasing randomness of target node dynamics). Only enhancing local structural connectivity around target nodes significantly reduced the brain network sensitivity (Figure 4). In addition, the reduced network sensitivity was highly correlated with the average node degree of target brain regions, which verifies our hypothesis on the central role of the hubs in reducing the network sensitivity. In contrast, decreasing the local connectivity and increasing/decreasing the randomness of node dynamics gave rise to the opposite results because they made the target sites functionally more separated from other nodes, hampering a hub-dominated progressive synchronization growth. Notably, the results imply that despite the importance of hub nodes as effective target sites, the type of modulation is also crucial to converting ES into GS network. These computational modeling results provide a theoretical criterion for developing an effective brain modulation method to reduce the hypersensitivity in FM. Irrespective of brain modulation methods, for instance, pharmacologic, electric, and magnetic stimulation, the outcomes of brain modulation should avoid significant decreases in the hub connectivity and significant change in the hub dynamics (i.e., decreasing/increasing randomness of local brain activity around hubs) that can enhance the ES condition as well as the network sensitivity in the brain.

### Modulating the brain network at criticality and its relationship with fibromyalgia

We implemented criticality to the brain network model to simulate spontaneous brain activities in a conscious resting state and examined the network sensitivity change between the critical states of the networks before and after the modulation. Brain activities in the conscious resting state share various signal properties of a system near criticality: large variance, large autocorrelation, long-range correlation, high susceptibility, and power-law (Beggs & Plenz, 2003; H. Kim & Lee, 2019; Shew et al., 2011; Tagliazucchi et al., 2016). As far as we know, modulating the brain network at criticality and examining the dependence of the network sensitivity on different types of network modulation, especially for the brain network occurring ES, have not been studied yet. In addition, this computational modeling study and the empirical evidence of ES in FM brain give us insights into the potential application of other important characteristics of ES and its relationship to chronic pain. In general, a system with ES conditions can be characterized by a hysteresis phenomenon, that is, differential pathways between forward and backward state transitions (i.e. transitions from incoherent to synchronized states and vice versa) (Boccaletti et al., 2016; Gómez-Gardeñes et al., 2011). Hysteresis is a universal phenomenon observed in nature and has been investigated in various research fields such as physics, biology, and engineering, as well as state transitions in the brain such as sleep and general anesthesia (Bertotti & Mayergoyz, 2006; Chikazumi et al., 1997; Friedman et al., 2010; Joiner et al., 2013; Rempe et al., 2010; Steyn-Ross et al., 2004; “The Elastic Hysteresis of Steel,” 1912; Voss et al., 2012). In particular, it has been found in general anesthesia of diverse species (drosophila, murine, mouse, rat, and human) that the anesthetic concentration required for inducing unconsciousness is higher than the concentration where consciousness is regained during general anesthesia (Luppi et al., 2021; McKinstry-Wu et al., 2020; Sepúlveda et al., 2019). In addition, our previous study has demonstrated that human subjects who have larger ES conditions in electroencephalogram (EEG) networks show larger hysteresis in state transitions during the loss and recovery of consciousness (H. Kim et al., 2018), implying individuals with larger ES conditions take longer time to recover from general anesthesia. Therefore, we expect that the brain with larger ES conditions such as FM brain experiences a large hysteresis phenomenon during state transition, for instance, faster loss of consciousness and more prolonged recovery in anesthesia.

In addition, another representative characteristic of ES at a critical point is the high instability of functional connectivity, which is originated from the large variance of network synchronization at a critical point (M. Kim & Lee, 2020). In a GS network, the structural connectivity significantly shapes functional connectivity patterns at a critical point with a large correlation (H. Kim et al., 2018; M. Kim et al., 2017). We have mathematically proved the underlying mechanism of this large correlation, empirically confirmed it with brain networks of different species (mouse, monkey, and human), and suggested the large correlation between functional and structural connectivity as an indicator of brain criticality (H. Lee et al., 2019). However, ES network has an aberrant hub network organization because the hub connectivity of the ES network is directly or indirectly disrupted. Considering the essential role of hubs in the organization of higher-order brain functions such as cognition (Vatansever et al., 2020; S. Wang et al., 2021; Zdanovskis et al., 2020), such aberrant functional brain organization may be associated with the cognitive deficit in FM. However, the detailed association between the cognitive deficit and the aberrant hub structure in the brain remains to be tested.

### Implication for effective brain modulation in fibromyalgia

The identified effective target sites in the human brain network model with the ES condition provide many implications for developing an effective brain modulation method for FM. The simulation results showed that increasing the local structural connectivity centered on hub regions including the insula, left isthmus cingulate cortex, and right precuneus is the most effective way to reduce the brain network sensitivity changing ES to GS. Interestingly, the brain regions such as the insula and the cingulate cortex have been already known as hubs in FM brain and currently take into account as the target sites for brain modulation to reduce the pain intensity (Kaplan et al., 2020). Recently, the motor cortex (M1) has been also considered as a target site for transcranial direct current stimulation (tDCS) for pain treatment (Cummiford et al., 2016; Foerster et al., 2015). Here we suggest the right precuneus, which is a hub region in the cortex, as a feasible target site for non-invasive brain stimulation such as tDCS to treat FM because of the limitation of direct stimulation to subcortical regions.

### Limitation

This study contains several limitations. First, the Stuart-Landau model has limitations in interpreting the simulation results because the model is not based on a biophysical mechanism. Instead of simulating the realistic brain activities, we focused on identifying a general network principle that could be associated with the emergence of hypersensitivity in the brain networks and effectively reduce the hypersensitivity at a global brain network level. This general network principle is applicable to diverse brain networks with different genders, ages, and diseases. Second, the target brain sites that we suggested to reduce the brain network sensitivity were selected from an anatomical brain network structure averaged over individuals. Thus, acquiring individual brain network structures would enable us to identify individualized target sites that might result in better performance to convert ES into GS. Third, the modulation we induced can reflect neuroplasticity and cortical excitability changes that are empirically observed in tDCS trials but cannot explicitly show it changes the excitatory or inhibitory connectivity. Fourth, the effects of the four types of brain network modulation are not independent of each other. However, despite the theoretic limitation of separating the intertwined effects, the different types of brain modulation (increasing/decreasing local connectivity or randomness of local node dynamics) provide us measurable aims that can be achieved with the experimental design. We expect that future studies with a fine model setting can address some of these limitations.

## Conclusion

This computational model study suggests a network mechanism-based brain modulation method that can significantly reduce brain network sensitivity. Increasing the local structural connectivity centered on hubs such as insular, isthmus cingulate cortex, and precuneus fundamentally changes the type of state transition from a 1^st^-order synchronization transition (ES) to a 2^nd^-order synchronization transition (GS), mitigates the ES condition, and reduces network sensitivity of the brain. The ES mechanism-based brain network modulation will provide a theoretic framework to design an effective brain stimulation method to systematically control the hypersensitivity in chronic pain patients.

## Materials and methods

The whole computational modeling procedures were summarized as the following (See Figure 1 for a schematic diagram of the study design).

1. We constructed a large-scale human brain network model containing ES as in the FM brain. We assumed that the large network sensitivity of ES is the underlying mechanism of hypersensitivity of FM brain, and the degree of ES condition correlates with the pain intensity of FM patients according to our previous empirical study (U. Lee et al., 2018).
2. We then modulated the degree of ES condition in the human brain network model with four different types of network modulation: increasing/decreasing the local structural connectivity and increasing/decreasing the randomness of node dynamics within a certain diameter centered on a target node.
3. The critical states for the brain network models were identified by searching through parameter space, at which the simulated brain activities reflect the characteristic brain dynamics during a conscious resting state. The brain networks with different ES and GS conditions had their own distinctive critical states.
4. We evaluated the ES condition (correlation between node degrees and frequencies) and the change of brain network sensitivity before and after a brain network modulation. We directly induced a global pulsatile stimulus to the brain network models before and after the modulations and quantified the changes of brain network sensitivity with surrogate measures— responsivity and Lempel Ziv Complexity.
5. Finally, we identified the type of network modulation and effective stimulation sites that can reduce the ES condition and the brain network sensitivity.

### Construction of ES human brain network model

We used a coupled Stuart-Landau model to simulate the oscillatory dynamics of the human brain network. The coupled Stuart-Landau model with an anatomically informed brain network structure has been widely applied to simulate the signals from various types of imaging modalities including EEG, magnetoencephalogram (MEG), and functional magnetic resonance imaging (fMRI) (Cabral et al., 2014, 2017; Deco et al., 2017, 2018; H. Kim et al., 2018; Kim & Lee, 2019; Moon et al., 2015). Since the goal of this study is to convert ES brain network into GS brain network, we used a modified Stuart-Landau model including an adaptive feedback term. Our previous study introduced the model to simulate a hysteresis phenomenon during general anesthesia (H. Kim et al., 2018). With this model, we can adjust the ES strengths of the brain network by modulating the adaptive feedback term 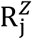, i.e., a recursive interaction process between a node j and the other nodes. The modified coupled Stuart-Landau model, composed of the *N* number of oscillators, is defined as the following:

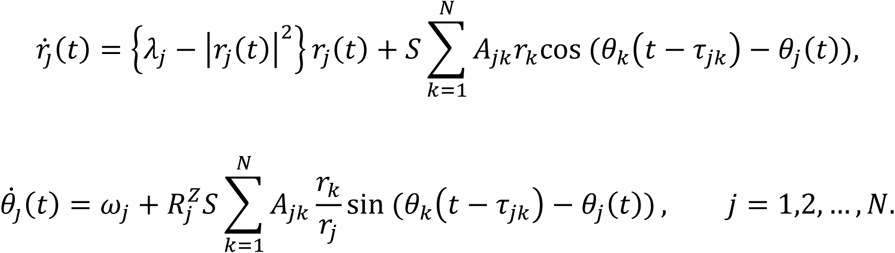

Here r_j_(*t*) is the amplitude of oscillator (node) j at time t. *λ*_j_ is a parameter modulating the randomness of the amplitude dynamics and S is a coupling strength among anatomically connected oscillatory dynamics. Changing these parameters induces various amplitude and phase dynamics through competition of independent node dynamics and its topological connections with other oscillators upon an anatomical brain network structure (Cavanna et al., 2018). Each node shows supercritical Hopf bifurcation, and the dynamics of the oscillator settle on a limit cycle if *λ*_*j*_ > 0, and on a stable fixed point if *λ*_*j*_ < 0. A_jk_ is the anatomical connection weight between oscillator j and k. The connection matrix A consists of 82 brain regions including cortical and subcortical regions which were constructed from diffusion tensor imaging (DTI) (van den Heuvel & Sporns, 2011). A certain value from 0 to 1 was assigned to A_jk_ based on the white matter connection weight between brain region j and k. The τ_jk_ = D_jk_/s, is a time delay between region j and k, where D_jk_ is a distance between brain regions with *s* = 7 ms, an average speed of axons across brain regions (Caminiti et al., 2013). The node *j* interacts with a connected node *k* after the time delay *τ*_*jk*_. θ_j_(*t*) is a phase of the oscillator j at time t, and ω_j_ is a natural frequency of the oscillator j. *R*_*j*_ is defined as 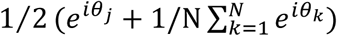, which is the extent of synchrony of node j with the other nodes. The *R*_*j*_ enhances a heterogeneity in the amplitude and phase dynamics of the brain network, incorporating the memory of a given synchronized state into the dynamics. The Z is a scale term for the adaptive feedback process *R*^*j*^, and it plays a role of a control parameter for adjusting ES strengths of the brain network model, indicating it inhibits a giant synchronization cluster formation centered around a node j with large synchrony, which is a core mechanism of ES (H. Kim et al., 2018).

We used Gaussian distribution for the natural frequency *ω* with a mean frequency of 10 Hz and a standard deviation of 0.4 Hz to simulate the dominant frequency bandwidth of human EEG alpha activity (H. Kim et al., 2018; M. Kim et al., 2017; U. Lee et al., 2018; Moon et al., 2015, 2017). We first fixed *λ*_*j*_ ≡ *λ* as 0. The coupling strength among the oscillators *S* was modulated from 0 to 30 with *δS* = 0.2, yielding the change of brain network from a fully incoherent state to a fully synchronized state. We set *Z* = 3, which is large enough to simulate ES in the brain network. We numerically solved differential equations of the Stuart-Landau model using the Runge-Kutta 4th method with 1,000 discretization steps and resampled the data with 250 Hz. For every coupling strength, the last 40 seconds of spontaneous oscillatory dynamics were used for the analysis after 20 seconds of saturation periods. Thirty different initial frequency configurations were simulated in each parameter set, and the results were averaged over all configurations.

### Four types of brain network modulation to convert ES into GS network

According to our hypothesis, enhancing hub connectivity between brain network nodes may facilitate giant synchronization cluster formation, which in turn converts ES network into GS network, reducing the brain network sensitivity significantly. In addition, it is known that repetitive tDCS targeting a local brain region can induce a long-lasting change in the brain network activity through neuroplasticity and cortical excitability shifts (Nitsche et al., 2007). Therefore, we tested four types of brain network modulation: increasing/decreasing the regional structural connectivity centered on a target node and increasing/decreasing the randomness of the regional activities centered on a target node in the ES human brain network model. These types of modulation may reflect neuroplasticity and cortical excitability shifts that are empirically observed in tDCS experiments. For changing the regional connectivity, we increased or decreased the local structural connectivity by three times within a radius of 2 cm centered on the targeted brain region, roughly simulating the effective range for connectivity change covered by a tDCS patch. For changing the randomness of regional brain activities, we increased or decreased the bifurcation parameter *λ* of nodes, respectively *λ* = 2 and *λ* = −2, within a radius of 2 cm centered on the target brain region. The parameters were chosen for simulating the most realistic results after the parameter test. The 30 highest-degree nodes in the human brain network were selected as the target brain regions and tested the modulation effect for each target brain region (Left and right putamen, amygdala, pallidum, thalamus, hippocampus, caudate, temporal pole, insula, precuneus, entorhinal cortex, precentral cortex, superior parietal cortex, accumbens-area, isthmus cingulate cortex, left superior frontal cortex, left postcentral cortex). Extreme cases such as *λ* = 5 or *λ* = −5 were presented in Supporting information Figure S8. The initial frequency configurations of the brain network models are all the same for the four types of modulation.

### Identification of critical states for the brain network model

Recent computational modeling and empirical studies suggest that brain dynamics in conscious states are near criticality (i.e., an optimally balanced state between order and disorder), and losing criticality is related to altered states of consciousness (Haimovici et al., 2013; H. Kim & Lee, 2019; Kitzbichler et al., 2009; Muñoz, 2018; Tagliazucchi et al., 2012). The network dynamics at a critical state reflect the characteristics of the brain activities in the conscious resting state: an optimal balance between stability and instability, optimal information processing, large flexibility to adapt to a changing environment, and wide repertoires of brain states (Beggs, 2008; Cocchi et al., 2017; Haimovici et al., 2013; H. Kim & Lee, 2019; Kitzbichler et al., 2009; Tagliazucchi et al., 2012). Therefore, we assumed that the critical states of a brain network before and after modulation of ES condition might reflect the distinctive brain activities in conscious states before and after the brain modulation. In this modeling study, we searched the parameter space of various coupling strengths and bifurcation parameters and determined the parameter set having a maximal autocorrelation function (ACF) as a critical state. The maximal ACF is one of the characteristics of a system approaching a critical transition, which is called “critical slowing down” and refers to the tendency of a system to take longer to return to equilibrium after a perturbation (Scheffer et al., 2009). To calculate the ACF of a simulated brain signal, we selected the time lag of 50-msec, which catches the dynamics of the alpha oscillations in the simulated brain signals.

### Evaluation of brain network sensitivity for the four types of brain network modulation

To test which type of brain modulation effectively reduces the brain network sensitivity and changes the type of state transition from ES (1^st^ order phase transition) to GS (2^nd^order phase transition), we investigated three different properties. (1) We examined the typical phase transitions of ES and GS. Converting ES into GS should present the typical change in transition pattern from a discrete synchronization transition (1^st^ order phase transition) to a continuous synchronization transition (2^nd^ order phase transition). (2) We examined whether a brain network modulation induces the typical network configuration of ES, a positive correlation between node degrees and node frequencies in the brain network. (3) Then, we directly measured the change of network sensitivity after the modulation directly implementing an external stimulus to the brain network.

To investigate the shape of synchronization transition, the instantaneous network synchronization *r*(*t*) at time *t* was measured by the order parameter of the oscillators,

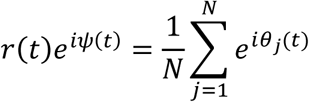

where *θ*_*j*_(*t*) is a phase of *j*_*th*_ oscillator and *ψ*(*t*) is the average global phase at time *t*. Here *r*(*t*) equals 0 if phases of oscillators are uniformly distributed and 1 if all oscillators have the same phase. The global network synchronization *R* is calculated by taking the average of the instantaneous network synchronization over time. As the coupling strengths increase in the brain networks before and after the four types of modulation, we investigated the shape of R to determine whether the transition is discrete or continuous.

A positive correlation between node degrees and frequencies was introduced as one of the network configurations that can induce ES in a heterogeneous network (Gómez-Gardeñes et al., 2011). Therefore, we tested which type of brain network modulation and which target node effectively mitigate the ES condition. The degree of *j*_*th*_ node was calculated by 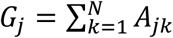 and the frequency of the *j*_*th*_ node was calculated using the instantaneous phases of the *j*_*th*_ node in the brain network at a critical state. A Spearman correlation between the degrees and frequencies, *ρ*^*deg*−*freq*^, was calculated for all the brain networks before and after modulation of different target nodes.

Finally, we directly measured the network sensitivity against an external stimulus and investigated which type of brain network modulation significantly reduces the brain network sensitivity. For the brain networks before and after the four types of modulation, we induced a global pulsatile stimulus to the brain networks and quantified the change of the network sensitivity. The global pulsatile stimulus was induced with *u*(*t*) as the following:

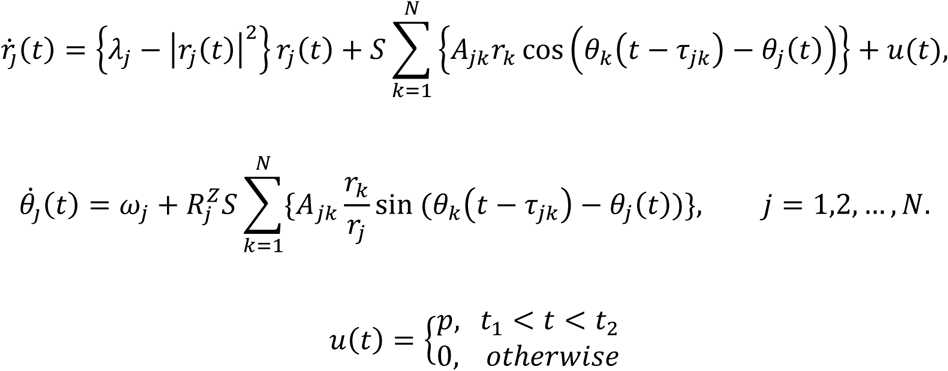

Here, *p* is the intensity of the stimuli during a period *T* = *t*_2_ − *t*_1_. We fixed *p* = 20 and duration *T* = 100 ms based on our previous findings (M. Kim and U. Lee 2020). The global stimulus was given at a randomly selected timing *t*_1_ for each trial. Twenty different trials were performed for each thirty different initial frequency configurations. Therefore, the same external stimulus was given 600 times to all the brain networks before and after modulation.

The sensitivity was obtained by measuring the responsivity and Lempel-Ziv complexity of the brain network’s response after the stimulation (M. Kim and U. Lee 2020). To measure the responsivity and complexity, we first calculated an instantaneous amplitude of the nodes for each stimulation trial. The instantaneous amplitude value for the *j*_*th*_ node of one trial was normalized by the mean and standard deviation of the baseline amplitude values of the *j*_*th*_ node. The baseline amplitude values were obtained from a total of 200-sec, consisting of 20 different stimulation trials of 10-sec pre-stimulus segment for each trial. We considered (1 − *α*) * 100^*th*^ percentile with *α* = 0.05 as a significantly changed amplitude after a stimulus onset. A perturbation response (PR) of the amplitude for the *j*_*th*_ node at time *t* was defined in a binary fashion: *PR*_*j*_(*t*) = 1, if the amplitude after the stimulus onset is significantly changed, and *PR*_*j*_(*t*) = 0, otherwise.

The responsivity *P* and Lempel-Ziv complexity *LZc* were calculated with the binary PR during 300 msec after the stimulus onset. The responsivity *P* was calculated by taking the average of *PR*(*t*) for all nodes over 300 msec. The *LZc*(*t*) of the *PR*(*t*) was defined as below:

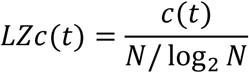

Here *c*(*t*) is the nonnormalized Lempel-Ziv complexity calculated by LZc76-algorithm [Lempel & Ziv, 1976], which is the number of unique patterns in the *PR*(*t*) at time *t*, and *N* is the number of nodes. The *LZc*(*t*) is the normalized *c*(*t*). The *LZc* was calculated by taking the average of *LZc*(*t*) over 300 msec after the stimulus onset. The *P* and *LZc* were calculated for each stimulation trial. The sensitivity results for each target brain regional modulation were the average of the 30 frequency configurations with the 20 stimulation trials.

## Acknowledgments

U.L., R.E.H., and A.F.D., were supported by National Institute of Health (NIH) grant R01AT010060. The funders had no role in study design, data collection and analysis, decision to publish, or preparation of the manuscript.

## Notes

### Competing Interest Statement

The authors have declared no competing interest.

## References

Beggs, J. M. (2008). The criticality hypothesis: How local cortical networks might optimize information processing. Philosophical Transactions of the Royal Society A: Mathematical, Physical and Engineering Sciences, 366(1864), 329–343. https://doi.org/10.1098/rsta.2007.2092

Beggs, J. M., & Plenz, D. (2003). Neuronal Avalanches in Neocortical Circuits. The Journal of Neuroscience, 23(35), 11167–11177. https://doi.org/10.1523/JNEUROSCI.23-35-11167.2003

Bertotti, G., & Mayergoyz, I. D. (2006). The science of hysteresis. Academic Press. https://search.ebscohost.com/login.aspx?direct=true&scope=site&db=nlebk&db=nlabk&AN=249064

Boccaletti, S., Almendral, J. A., Guan, S., Leyva, I., Liu, Z., Sendiña-Nadal, I., Wang, Z., & Zou, Y. (2016). Explosive transitions in complex networks’ structure and dynamics: Percolation and synchronization. Physics Reports, 660, 1–94. https://doi.org/10.1016/j.physrep.2016.10.004

Cabral, J., Kringelbach, M. L., & Deco, G. (2017). Functional connectivity dynamically evolves on multiple time-scales over a static structural connectome: Models and mechanisms. NeuroImage, 160, 84–96. https://doi.org/10.1016/j.neuroimage.2017.03.045

Cabral, J., Luckhoo, H., Woolrich, M., Joensson, M., Mohseni, H., Baker, A., Kringelbach, M. L., & Deco, G. (2014). Exploring mechanisms of spontaneous functional connectivity in MEG: How delayed network interactions lead to structured amplitude envelopes of band-pass filtered oscillations. NeuroImage, 90, 423–435. https://doi.org/10.1016/j.neuroimage.2013.11.047

Caminiti, R., Carducci, F., Piervincenzi, C., Battaglia-Mayer, A., Confalone, G., Visco-Comandini, F., Pantano, P., & Innocenti, G. M. (2013). Diameter, Length, Speed, and Conduction Delay of Callosal Axons in Macaque Monkeys and Humans: Comparing Data from Histology and Magnetic Resonance Imaging Diffusion Tractography. Journal of Neuroscience, 33(36), 14501–14511. https://doi.org/10.1523/JNEUROSCI.0761-13.2013

Cavanna, F., Vilas, M. G., Palmucci, M., & Tagliazucchi, E. (2018). Dynamic functional connectivity and brain metastability during altered states of consciousness. NeuroImage, 180, 383–395. https://doi.org/10.1016/j.neuroimage.2017.09.065

Chen, H., He, G., Huang, F., Shen, C., & Hou, Z. (2013). Explosive synchronization transitions in complex neural networks. Chaos (Woodbury, N.Y.), 23(3), 033124. https://doi.org/10.1063/1.4818543

Chikazumi, S., Graham, C. D., & Chikazumi, S. (1997). Physics of ferromagnetism (2nd ed). Clarendon Press; Oxford University Press.

Cocchi, L., Gollo, L. L., Zalesky, A., & Breakspear, M. (2017). Criticality in the brain: A synthesis of neurobiology, models and cognition. Progress in Neurobiology, 158, 132–152. https://doi.org/10.1016/J.PNEUROBIO.2017.07.002

Cummiford, C. M., Nascimento, T. D., Foerster, B. R., Clauw, D. J., Zubieta, J.-K., Harris, R. E., & DaSilva, A. F. (2016). Changes in resting state functional connectivity after repetitive transcranial direct current stimulation applied to motor cortex in fibromyalgia patients. Arthritis Research & Therapy, 18, 40. https://doi.org/10.1186/s13075-016-0934-0

Dai, X., Li, X., Gutiérrez, R., Guo, H., Jia, D., Perc, M., Manshour, P., Wang, Z., & Boccaletti, S. (2020). Explosive synchronization in populations of cooperative and competitive oscillators. Chaos, Solitons & Fractals, 132, 109589. https://doi.org/10.1016/j.chaos.2019.109589

Deco, G., Cabral, J., Saenger, V. M., Boly, M., Tagliazucchi, E., Laufs, H., Van Someren, E., Jobst, B., Stevner, A., & Kringelbach, M. L. (2018). Perturbation of whole-brain dynamics in silico reveals mechanistic differences between brain states. NeuroImage, 169, 46–56. https://doi.org/10.1016/j.neuroimage.2017.12.009

Deco, G., Kringelbach, M. L., Jirsa, V. K., & Ritter, P. (2017). The dynamics of resting fluctuations in the brain: Metastability and its dynamical cortical core. Scientific Reports, 7(1), 3095. https://doi.org/10.1038/s41598-017-03073-5

Foerster, B. R., Nascimento, T. D., DeBoer, M., Bender, M. A., Rice, I. C., Truong, D. Q., Bikson, M., Clauw, D. J., Zubieta, J.-K., Harris, R. E., & DaSilva, A. F. (2015). Excitatory and inhibitory brain metabolites as targets of motor cortex transcranial direct current stimulation therapy and predictors of its efficacy in fibromyalgia. Arthritis & Rheumatology (Hoboken, N.J.), 67(2), 576–581. https://doi.org/10.1002/art.38945

Friedman, E. B., Sun, Y., Moore, J. T., Hung, H.-T., Meng, Q. C., Perera, P., Joiner, W. J., Thomas, S. A., Eckenhoff, R. G., Sehgal, A., & Kelz, M. B. (2010). A conserved behavioral state barrier impedes transitions between anesthetic-induced unconsciousness and wakefulness: Evidence for neural inertia. PloS One, 5(7), e11903. https://doi.org/10.1371/journal.pone.0011903

Gollo, L. L., Zalesky, A., Hutchison, R. M., van den Heuvel, M., & Breakspear, M. (2015). Dwelling quietly in the rich club: Brain network determinants of slow cortical fluctuations. Philosophical Transactions of the Royal Society of London. Series B, Biological Sciences, 370(1668), 20140165. https://doi.org/10.1098/rstb.2014.0165

Gómez-Gardeñes, J., Gómez, S., Arenas, A., & Moreno, Y. (2011). Explosive synchronization transitions in scale-free networks. Physical Review Letters, 106(12), 128701. https://doi.org/10.1103/PhysRevLett.106.128701

Gómez-Gardeñes, J., Moreno, Y., & Arenas, A. (2007). Paths to Synchronization on Complex Networks. Physical Review Letters, 98(3), 034101. https://doi.org/10.1103/PhysRevLett.98.034101

Gu, S., Pasqualetti, F., Cieslak, M., Telesford, Q. K., Yu, A. B., Kahn, A. E., Medaglia, J. D., Vettel, J. M., Miller, M. B., Grafton, S. T., & Bassett, D. S. (2015). Controllability of structural brain networks. Nature Communications, 6, 8414. https://doi.org/10.1038/ncomms9414

Haimovici, A., Tagliazucchi, E., Balenzuela, P., & Chialvo, D. R. (2013). Brain organization into resting state networks emerges at criticality on a model of the human connectome. Physical Review Letters, 110(17), 178101. https://doi.org/10.1103/PhysRevLett.110.178101

Harris, R. E., Napadow, V., Huggins, J. P., Pauer, L., Kim, J., Hampson, J., Sundgren, P. C., Foerster, B., Petrou, M., Schmidt-Wilcke, T., & Clauw, D. J. (2013). Pregabalin rectifies aberrant brain chemistry, connectivity, and functional response in chronic pain patients. Anesthesiology, 119(6), 1453–1464. https://doi.org/10.1097/ALN.0000000000000017

Harte, S. E., Harris, R. E., & Clauw, D. J. (2018). The neurobiology of central sensitization. Journal of Applied Biobehavioral Research, 23(2). https://doi.org/10.1111/jabr.12137

Hawkins, R. A. (2013). Fibromyalgia: A clinical update. The Journal of the American Osteopathic Association, 113(9), 680–689. https://doi.org/10.7556/jaoa.2013.034

Honey, C. J., Kotter, R., Breakspear, M., & Sporns, O. (2007). Network structure of cerebral cortex shapes functional connectivity on multiple time scales. Proceedings of the National Academy of Sciences, 104(24), 10240–10245. https://doi.org/10.1073/pnas.0701519104

Joiner, W. J., Friedman, E. B., Hung, H.-T., Koh, K., Sowcik, M., Sehgal, A., & Kelz, M. B. (2013). Genetic and anatomical basis of the barrier separating wakefulness and anesthetic-induced unresponsiveness. PLoS Genetics, 9(9), e1003605. https://doi.org/10.1371/journal.pgen.1003605

Kaplan, C. M., Harris, R. E., Lee, U., DaSilva, A. F., Mashour, G. A., & Harte, S. E. (2020). Targeting network hubs with noninvasive brain stimulation in patients with fibromyalgia. Pain, 161(1), 43–46. https://doi.org/10.1097/j.pain.0000000000001696

Kaplan, C. M., Schrepf, A., Vatansever, D., Larkin, T. E., Mawla, I., Ichesco, E., Kochlefl, L., Harte, S. E., Clauw, D. J., Mashour, G. A., & Harris, R. E. (2019). Functional and neurochemical disruptions of brain hub topology in chronic pain. Pain, 160(4), 973–983. https://doi.org/10.1097/j.pain.0000000000001480

Kim, H., & Lee, U. (2019). Criticality as a determinant of integrated information Φ in human brain networks. Entropy, 21(10), 981. https://doi.org/10.3390/e21100981

Kim, H., Moon, J., Mashour, G. A., & Lee, U. (2018). Mechanisms of hysteresis in human brain networks during transitions of consciousness and unconsciousness: Theoretical principles and empirical evidence. 1–22.

Kim & Lee. (2019). Criticality as a Determinant of Integrated Information Φ in Human Brain Networks. Entropy, 21(10), 981. https://doi.org/10.3390/e21100981

Kim, M., Kim, S., Mashour, G. A., & Lee, U. (2017). Relationship of Topology, Multiscale Phase Synchronization, and State Transitions in Human Brain Networks. Frontiers in Computational Neuroscience, 11(June), 1–12. https://doi.org/10.3389/fncom.2017.00055

Kim, M., & Lee, U. (2020). Alpha oscillation, criticality, and responsiveness in complex brain networks. Network Neuroscience, 4(1), 155–173. https://doi.org/10.1162/netn_a_00113

Kim, M., Mashour, G. A., Moraes, S.-B., Vanini, G., Tarnal, V., Janke, E., Hudetz, A. G., & Lee, U. (2016). Functional and Topological Conditions for Explosive Synchronization Develop in Human Brain Networks with the Onset of Anesthetic-Induced Unconsciousness. Frontiers in Computational Neuroscience, 10. https://doi.org/10.3389/fncom.2016.00001

Kitzbichler, M. G., Smith, M. L., Christensen, S. R., & Bullmore, E. (2009). Broadband criticality of human brain network synchronization. PLoS Computational Biology, 5(3), 1000314. https://doi.org/10.1371/journal.pcbi.1000314

Lee, H., Golkowski, D., Jordan, D., Berger, S., Ilg, R., Lee, J., Mashour, G. A., Lee, U., & ReCCognition Study Group. (2019). Relationship of critical dynamics, functional connectivity, and states of consciousness in large-scale human brain networks. NeuroImage, 188, 228–238. https://doi.org/10.1016/j.neuroimage.2018.12.011

Lee, U., Kim, M., Lee, K., Kaplan, C. M., Clauw, D. J., Kim, S., Mashour, G. A., & Harris, R. E. (2018). Functional Brain Network Mechanism of Hypersensitivity in Chronic Pain. Scientific Reports. https://doi.org/10.1038/s41598-017-18657-4

Luppi, A. I., Spindler, L. R. B., Menon, D. K., & Stamatakis, E. A. (2021). The Inert Brain: Explaining Neural Inertia as Post-anaesthetic Sleep Inertia. Frontiers in Neuroscience, 15, 643871. https://doi.org/10.3389/fnins.2021.643871

McKinstry-Wu, A. R., Proekt, A., & Kelz, M. B. (2020). Neural Inertia: A Sticky Situation for Anesthesia. Journal of Neurosurgical Anesthesiology, 32(3), 190–192. https://doi.org/10.1097/ANA.0000000000000687

Menzies, V. (2016). Fibromyalgia Syndrome: Current Considerations in Symptom Management. The American Journal of Nursing, 116(1), 24–32; quiz 33, 41. https://doi.org/10.1097/01.NAJ.0000476162.13177.ae

Moon, J. Y., Kim, J., Ko, T. W., Kim, M., Iturria-Medina, Y., Choi, J. H., Lee, J., Mashour, G. A., & Lee, U. C. (2017). Structure shapes dynamics and directionality in diverse brain networks: Mathematical principles and empirical confirmation in three species. Scientific Reports. https://doi.org/10.1038/srep46606

Moon, J. Y., Lee, U. C., Blain-Moraes, S., & Mashour, G. A. (2015). General relationship of global topology, local dynamics, and directionality in large-scale brain networks. PLoS Computational Biology, 11(4), 1–21. https://doi.org/10.1371/journal.pcbi.1004225

Muñoz, M. A. (2018). Colloquium: Criticality and dynamical scaling in living systems. Reviews of Modern Physics, 90(3), 031001. https://doi.org/10.1103/RevModPhys.90.031001

Nitsche, M. A., Doemkes, S., Karaköse, T., Antal, A., Liebetanz, D., Lang, N., Tergau, F., & Paulus, W. (2007). Shaping the Effects of Transcranial Direct Current Stimulation of the Human Motor Cortex. Journal of Neurophysiology, 97(4), 3109–3117. https://doi.org/10.1152/jn.01312.2006

Rempe, M. J., Best, J., & Terman, D. (2010). A mathematical model of the sleep/wake cycle. Journal of Mathematical Biology, 60(5), 615–644. https://doi.org/10.1007/s00285-009-0276-5

Scheffer, M., Bascompte, J., Brock, W. A., Brovkin, V., Carpenter, S. R., Dakos, V., Held, H., van Nes, E. H., Rietkerk, M., & Sugihara, G. (2009). Early-warning signals for critical transitions. Nature, 461(7260), 53–59. https://doi.org/10.1038/nature08227

Schmidt, R., LaFleur, K. J. R., de Reus, M. A., van den Berg, L. H., & van den Heuvel, M. P. (2015). Kuramoto model simulation of neural hubs and dynamic synchrony in the human cerebral connectome. BMC Neuroscience, 16, 54. https://doi.org/10.1186/s12868-015-0193-z

Sepúlveda, P. O., Tapia, L. F., & Monsalves, S. (2019). Neural inertia and differences between loss of and recovery from consciousness during total intravenous anaesthesia: A narrative review. Anaesthesia, 74(6), 801–809. https://doi.org/10.1111/anae.14609

Shew, W. L., Yang, H., Yu, S., Roy, R., & Plenz, D. (2011). Information Capacity and Transmission Are Maximized in Balanced Cortical Networks with Neuronal Avalanches. Journal of Neuroscience, 31(1), 55–63. https://doi.org/10.1523/JNEUROSCI.4637-10.2011

Steyn-Ross, M. L., Steyn-Ross, D. A., & Sleigh, J. W. (2004). Modelling general anaesthesia as a first-order phase transition in the cortex. Progress in Biophysics and Molecular Biology, 85(2–3), 369–385. https://doi.org/10.1016/j.pbiomolbio.2004.02.001

Tagliazucchi, E., Balenzuela, P., Fraiman, D., & Chialvo, D. R. (2012). Criticality in large-scale brain fMRI dynamics unveiled by a novel point process analysis. Frontiers in Physiology, 3, 15. https://doi.org/10.3389/fphys.2012.00015

Tagliazucchi, E., Chialvo, D. R., Siniatchkin, M., Amico, E., Brichant, J.-F., Bonhomme, V., Noirhomme, Q., Laufs, H., & Laureys, S. (2016). Large-scale signatures of unconsciousness are consistent with a departure from critical dynamics. Journal of The Royal Society Interface, 13(114), 20151027. https://doi.org/10.1098/rsif.2015.1027

The elastic hysteresis of steel. (1912). Proceedings of the Royal Society of London. Series A, Containing Papers of a Mathematical and Physical Character, 87(598), 502–511. https://doi.org/10.1098/rspa.1912.0104

van den Heuvel, M. P., & Sporns, O. (2011). Rich-Club Organization of the Human Connectome. Journal of Neuroscience, 31(44), 15775–15786. https://doi.org/10.1523/JNEUROSCI.3539-11.2011

Vatansever, D., Schröter, M., Adapa, R. M., Bullmore, E. T., Menon, D. K., & Stamatakis, E. A. (2020). Reorganisation of Brain Hubs across Altered States of Consciousness. Scientific Reports, 10(1), 3402. https://doi.org/10.1038/s41598-020-60258-1

Vlasov, V., & Bifone, A. (2017). Hub-driven remote synchronization in brain networks. Scientific Reports, 7(1), 10403. https://doi.org/10.1038/s41598-017-09887-7

Voss, L. J., Brock, M., Carlsson, C., Steyn-Ross, A., Steyn-Ross, M., & Sleigh, J. W. (2012). Investigating paradoxical hysteresis effects in the mouse neocortical slice model. European Journal of Pharmacology, 675(1–3), 26–31. https://doi.org/10.1016/j.ejphar.2011.11.045

Wang, C.-Q., Pumir, A., Garnier, N. B., & Liu, Z.-H. (2017). Explosive synchronization enhances selectivity: Example of the cochlea. Frontiers of Physics, 12(5), 128901. https://doi.org/10.1007/s11467-016-0634-x

Wang, S., Gan, S., Yang, X., Li, T., Xiong, F., Jia, X., Sun, Y., Liu, J., Zhang, M., & Bai, L. (2021). Decoupling of Structural and Functional Connectivity in Hubs and Cognitive Impairment After Mild Traumatic Brain Injury. Brain Connectivity. https://doi.org/10.1089/brain.2020.0852

Zdanovskis, N., Platkājis, A., Kostiks, A., & Karelis, G. (2020). Structural Analysis of Brain Hub Region Volume and Cortical Thickness in Patients with Mild Cognitive Impairment and Dementia. Medicina (Kaunas, Lithuania), 56(10), E497. https://doi.org/10.3390/medicina56100497

Zhang, X., Guan, S., Zou, Y., Chen, X., & Liu, Z. (2016). Suppressing explosive synchronization by contrarians. EPL (Europhysics Letters), 113(2), 28005. https://doi.org/10.1209/0295-5075/113/28005

Zhang, X., Zou, Y., Boccaletti, S., & Liu, Z. (2015). Explosive synchronization as a process of explosive percolation in dynamical phase space. Scientific Reports, 4(1), 5200. https://doi.org/10.1038/srep05200

