## Supplementary materials for "Explosive Synchronization-Based Brain Modulation Reduces Hypersensitivity in The Brain Network: A Computational Model Study"

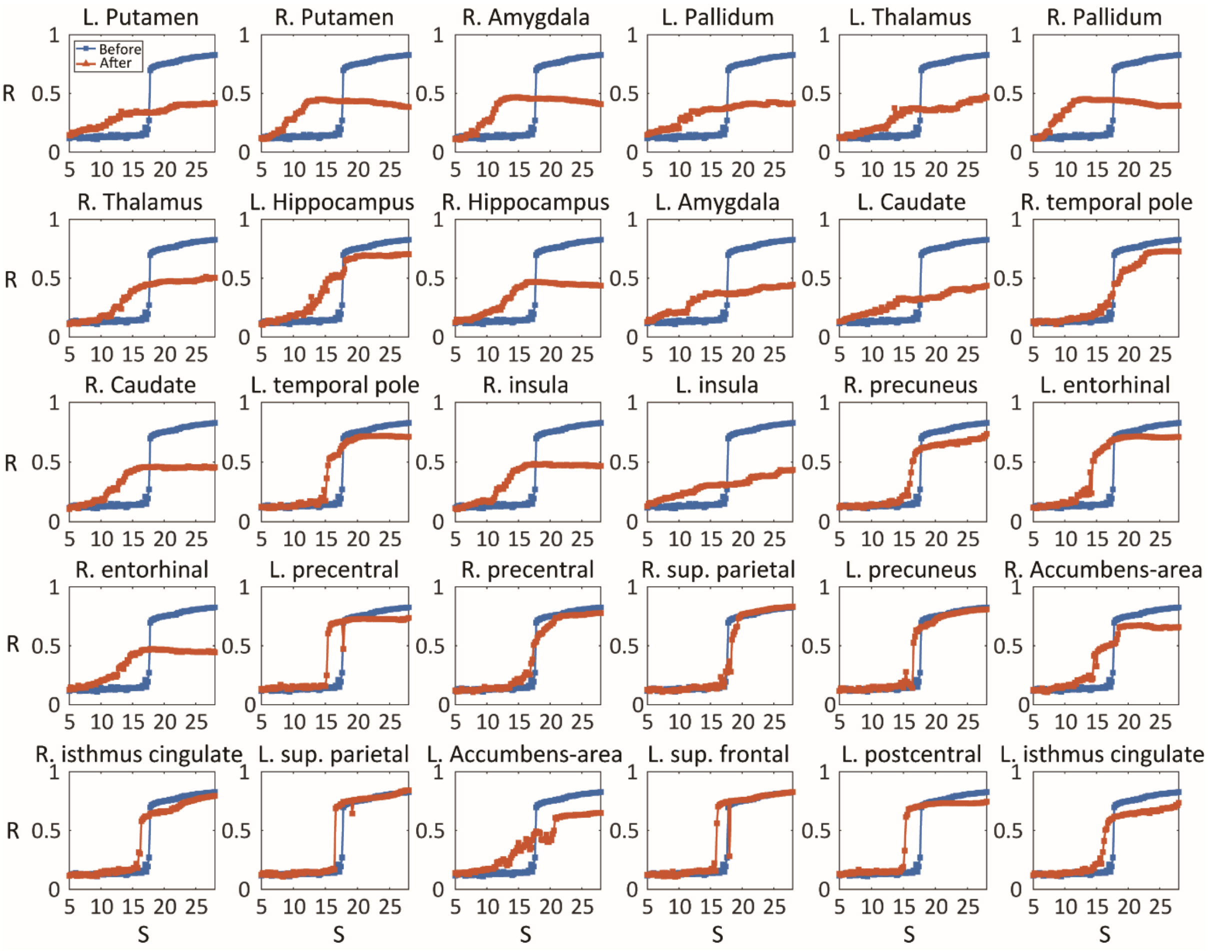


**Figure S1**. The synchronization transition shape for the node connectivity increase (CI) modulation. Each subplot shows the synchronization transition shape of brain networks before (blue) and after (orange) the modulation centered around a specific node. The modulated center node for each node modulation is shown on the title of the subplot. Most of the CI node modulation change the shape of transition from abrupt to gradual form.

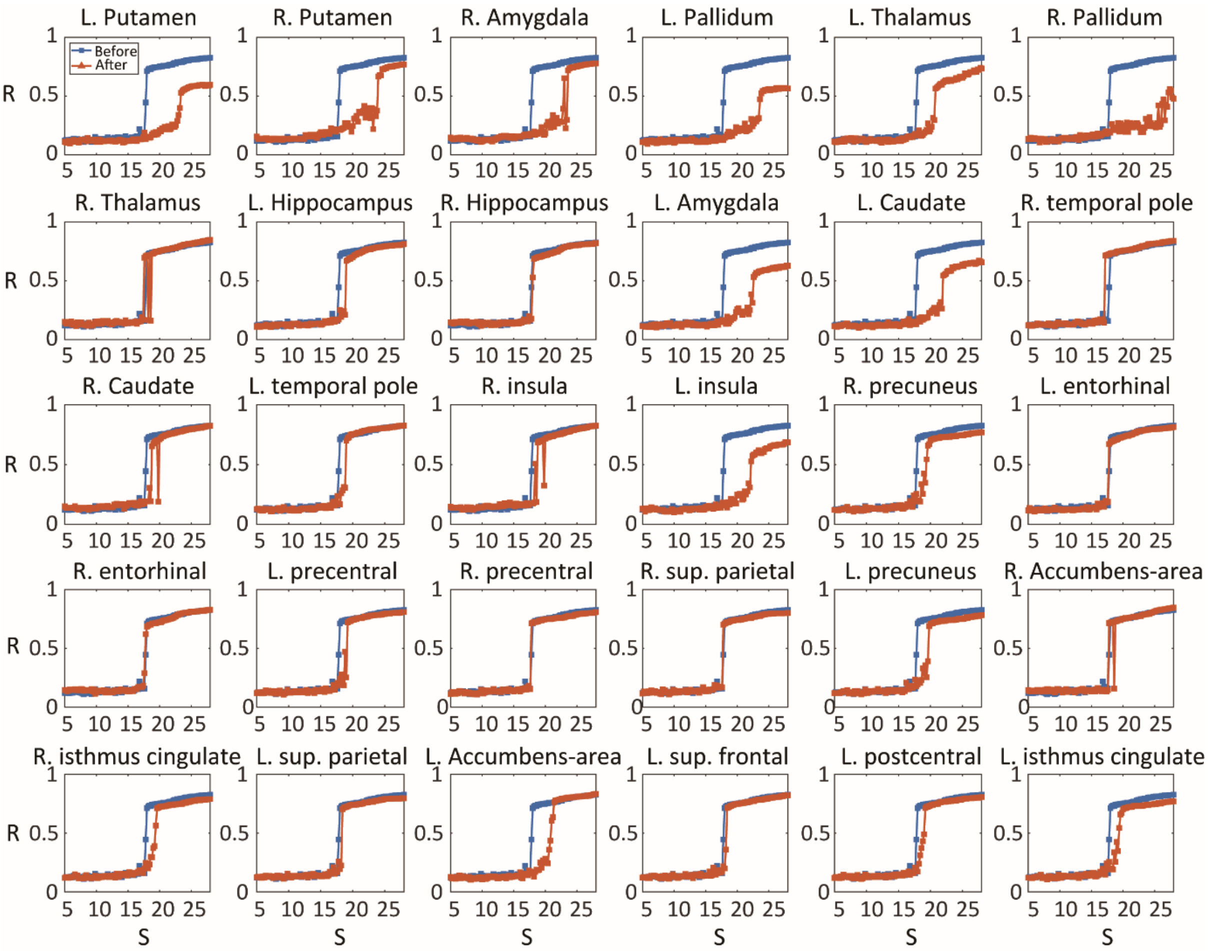


**Figure S2**. The synchronization transition shape for the node connectivity decrease (CD) modulation. Each subplot shows the synchronization transition shape of brain networks before (blue) and after (orange) the modulation centered around a specific node. The modulated center node for each node modulation is shown on the title of the subplot. Most of the CD node modulation cannot change the shape of transition.

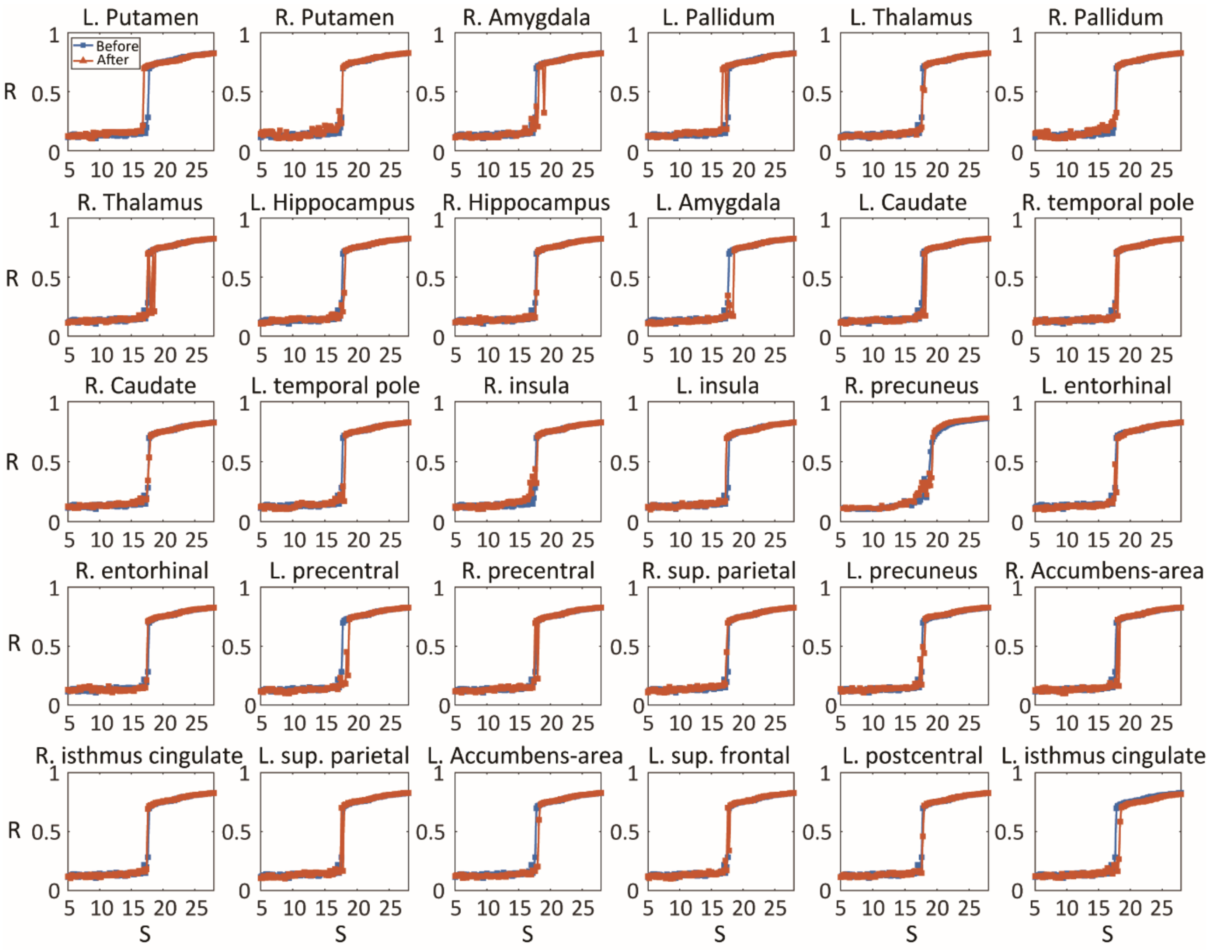


**Figure S3**. The synchronization transition shape for the node randomness increase (RI) modulation. Each subplot shows the synchronization transition shape of brain networks before (blue) and after (orange) the modulation centered around a specific node. The modulated center node for each node modulation is shown on the title of the subplot. Most of the RI node modulation cannot change the shape of transition.

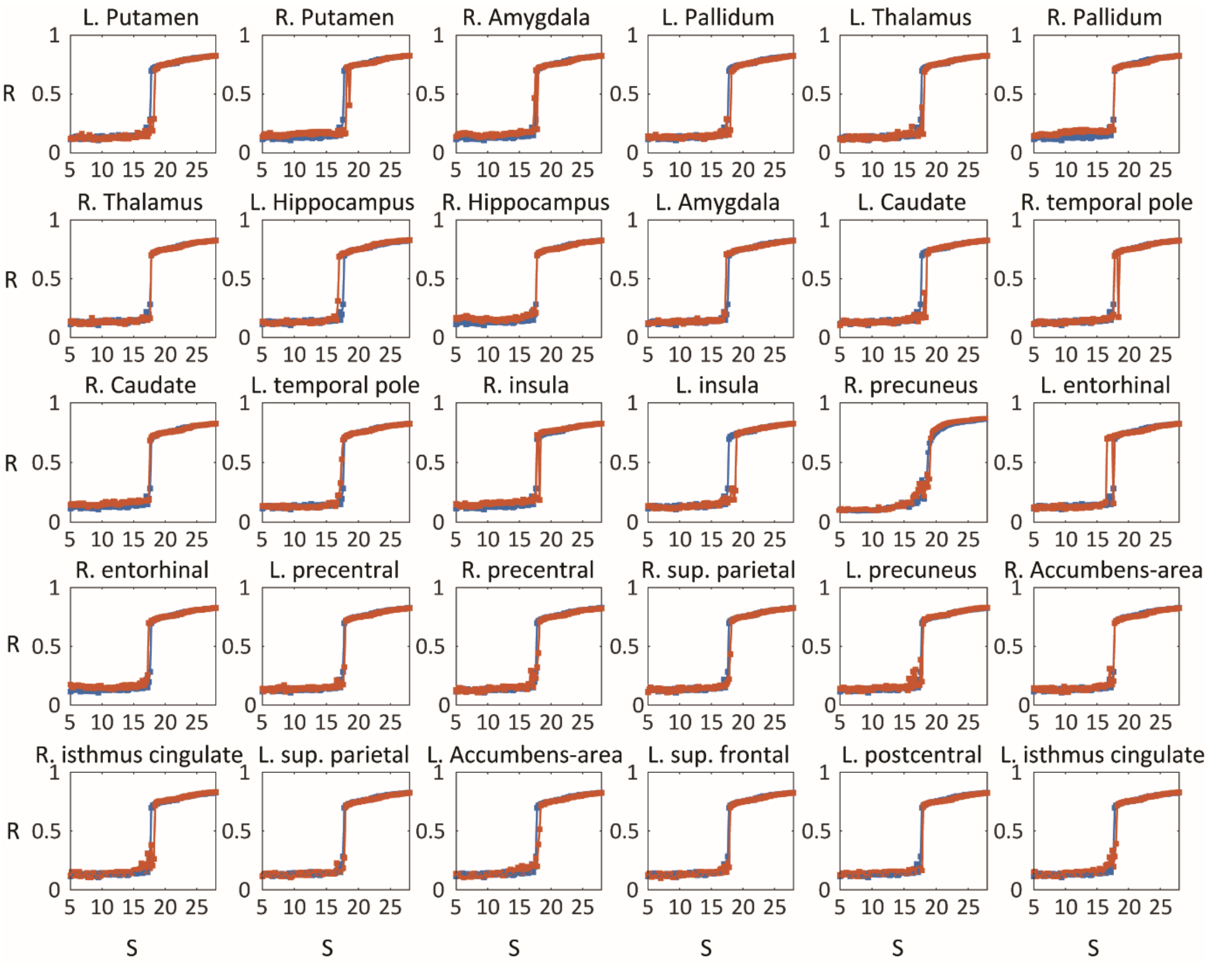


**Figure S4**. The synchronization transition shape for the node randomness decrease (RD) modulation. Each subplot shows the synchronization transition shape of brain networks before (blue) and after (orange) the modulation centered around a specific node. The modulated center node for each node modulation is shown on the title of the subplot. Most of the RD node modulation cannot change the shape of transition.

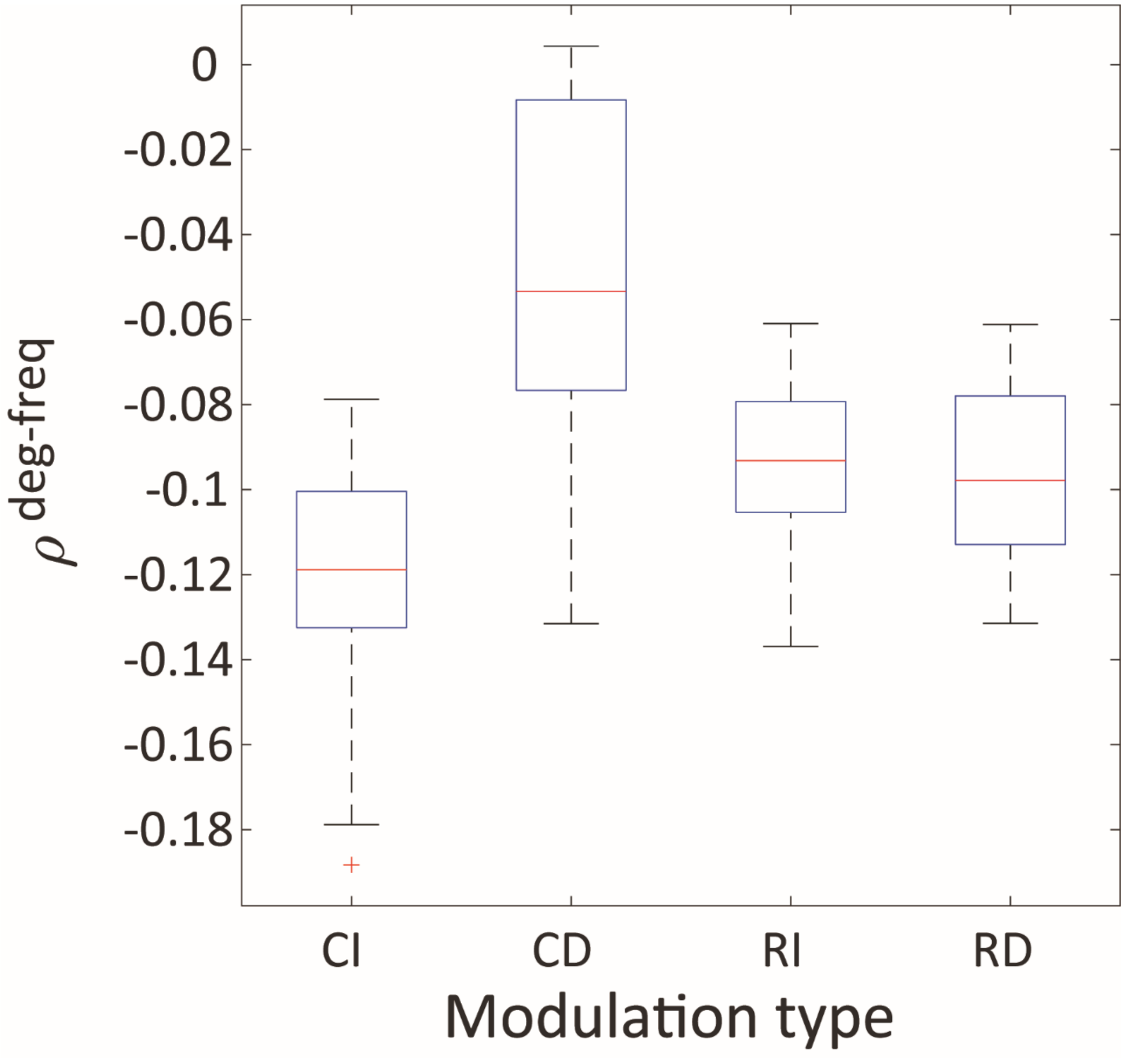


**Figure S5**. The average correlation between the node degree and frequency $\rho^{deg-freq}$ for four types of modulation (CI, CD, RI, and RD). On each box, the central red line indicates the median, and the bottom and top edges of the blue box indicate the 25th and 75th percentiles, respectively. The whiskers present the extreme data points. The red '+' symbol indicates the outliers. The correlation of CI modulation is significantly lower than the other types of modulation (Kuruskal-Wallis test with multiple comparison: $p<0.001$ for CI vs. CD; $p<0.01$ for CI vs. RI; $p<0.005$ for CI vs. RD)

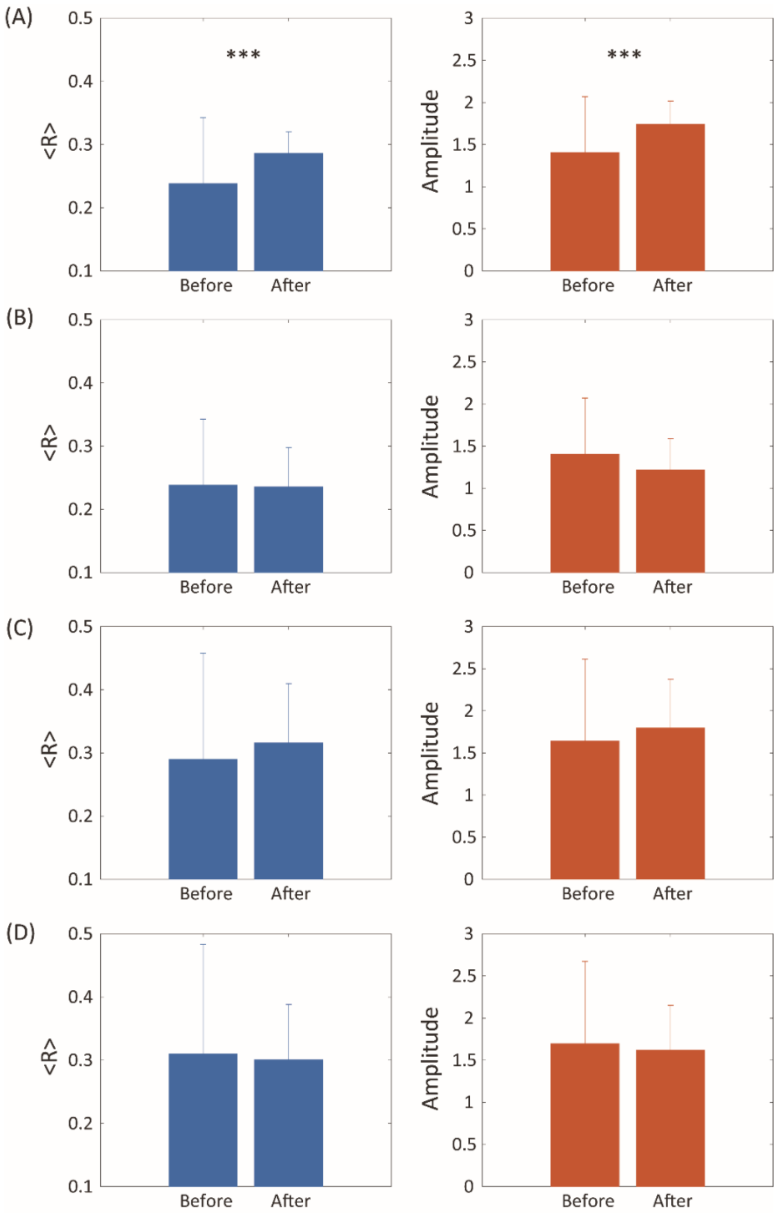


**Figure S6**. The average global synchronization <R> and a hub node amplitude of signals for four types of modulation: (A) CI, (B) CD, (C) RI, and (D) RD. The bar indicates an average value of <R> (blue) and an average value of hub node amplitude (orange) of thirty iterations for thirty different node modulations. The error bar indicates standard deviation of thirty different iterations. The <R> and node amplitude were significantly increased in the CI modulation (Wilcoxon Rank-sum test, $p<0.01$), but it was not significantly changed for other types of modulation.

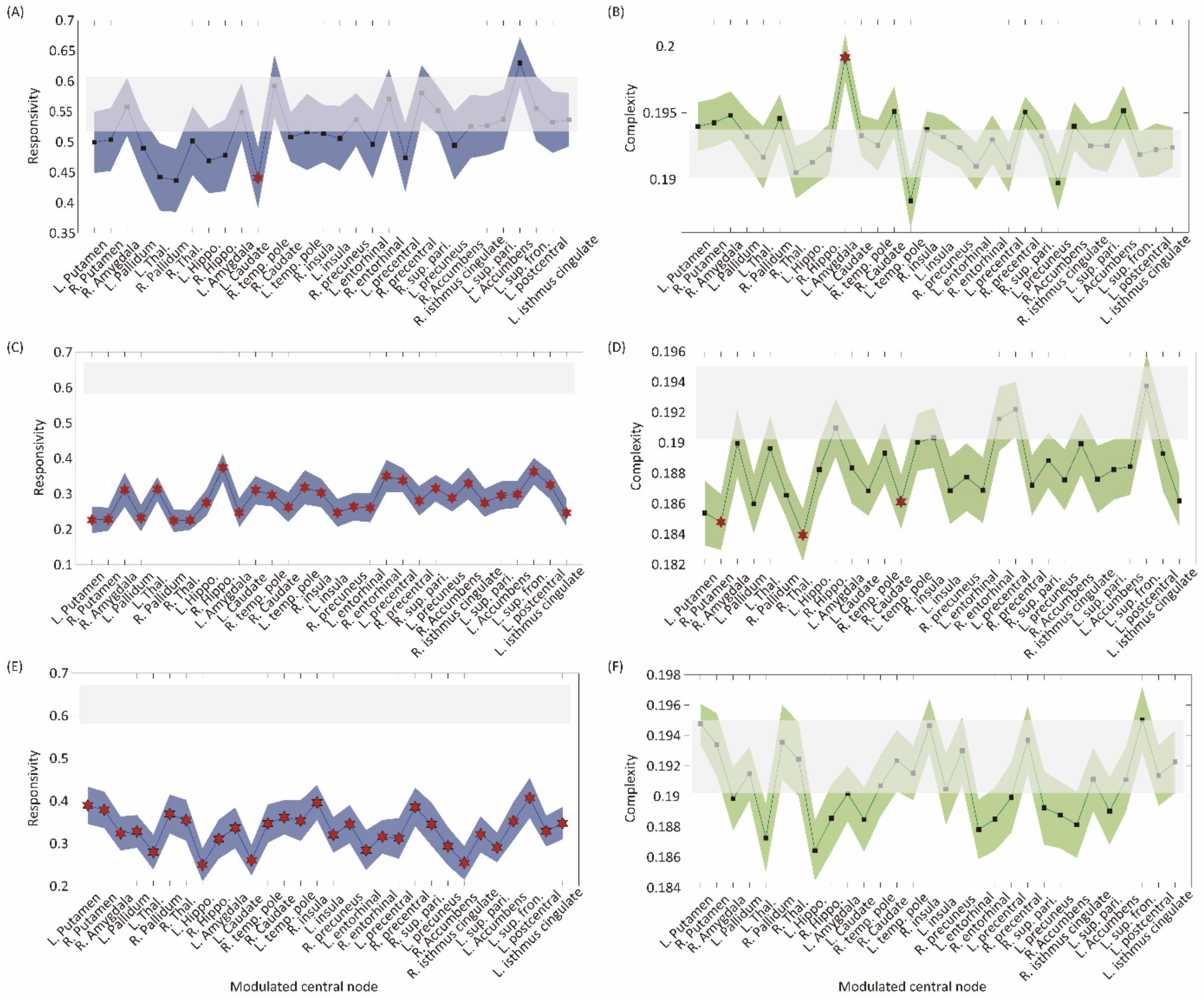

**Figure S7**. Brain network sensitivity to external stimuli in CD, RI, and RD modulations. Responsivity (blue) and complexity (green) of networks before and after the CD (A and B), RI (C and D), and RD (E and F) modulation are presented. The responsivity was decreased in RI and RD modulation, but the complexity of most of the node modulation was not significantly changed except for right putamen, right thalamus, and right caudate of RI modulation.

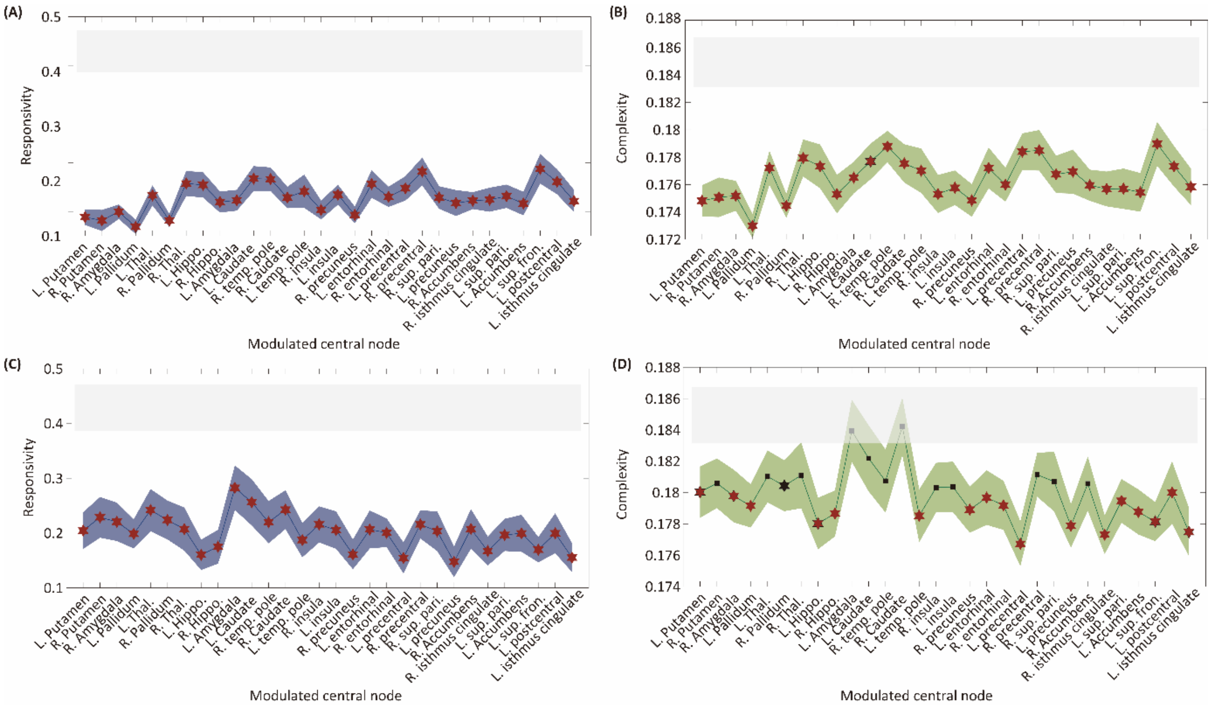


**Figure S8**. Brain network sensitivity to external stimuli for RI, and RD modulations with bifurcation parameters $\lambda = -5$ and $\lambda = 5$. Responsivity (blue) and complexity (green) of brain networks before and after the RI (A and B) and RD (C and D) modulation are presented. The responsivity and complexity were decreased in most of the node modulation of both RI and RD modulation.
